# Social isolation recruits amygdala-medial prefrontal cortex projections to escalate alcohol drinking in male mice

**DOI:** 10.1101/2023.11.09.566421

**Authors:** Reesha R. Patel, Kelly N. Kim, Makenzie Patarino, Rachelle Pamintuan, Felix H. Taschbach, Hao Li, Bitna Joo, Anna Pallé, Xianru Yu, Christopher R. Lee, Aniek van Hoek, Jesse White, Rogelio Castro, Christian Cazares, Raymundo L. Miranda, Caroline Jia, Jeremy Delahanty, Kanha Batra, Laurel R. Keyes, Avraham Libster, Romy Wichmann, Talmo D. Pereira, Marcus K. Benna, Kay M. Tye

**Author notes:** These authors contributed equally (Reesha R. Patel, Kelly N. Kim). Co-Corresponding Authors: Correspondence: Reesha R. Patel, Ph.D. Assistant Professor Feinberg School of Medicine, 320 E. Superior St., Chicago, IL 60612, USA @PatelReesha, Kay M. Tye, Ph.D. HHMI Investigator, Wylie Vale Professor, Salk Institute for Biological Studies 10010 North Torrey Pines Rd., La Jolla, CA 92037, USA, @kaymtye.

## Abstract

Social isolation profoundly alters motivation and increases vulnerability to alcohol misuse in humans, yet the underlying neural mechanisms remain unclear. Here we show that isolation escalates alcohol drinking in male mice but suppresses it in females. Whole-cell recordings revealed that neurons in the basolateral amygdala projecting to the medial prefrontal cortex (BLA-mPFC) track alcohol intake in both sexes. Isolation increased BLA-mPFC excitability in males but decreased it in females, mirroring their opposite behavioral adaptations. Given this divergence, we focused subsequent mechanistic studies on males to isolate neural pathway-level drivers of escalated alcohol intake. Cellular-resolution calcium imaging showed that activity in BLA-mPFC neurons encodes and predicts alcohol drinking, and optogenetic activation of this pathway increased alcohol intake. Simultaneous optogenetics and calcium imaging revealed that BLA-mPFC stimulation enhanced mPFC neuronal responses to alcohol, mimicking isolation-induced activity patterns, while photoinhibition reduced drinking in isolated mice. Together, these findings identify a BLA-mPFC pathway mechanism through which social isolation reconfigures prefrontal processing to promote alcohol intake.

## INTRODUCTION

Alcohol use disorder (AUD) is the most prevalent substance use disorder worldwide^1^, affecting an estimated 7% of the global population over age 15, according to the World Health Organization. While many factors influence the decision to drink alcohol, some have a greater propensity to predispose individuals to long-term detriments of misuse.

Social determinants, including social isolation, have been linked to increased alcohol consumption across species, from rodents to humans^2–8^, underscoring the conserved impact of social environment on alcohol use. Social isolation is emerging as an increasingly relevant risk factor in humans^9–11^, particularly in adulthood, when isolation is more common^10,12^ and exerts greater negative impact^13^. Consistent with this, social isolation in adults is associated with heavy drinking in humans^5^ and increased alcohol intake in mice^6,14–16^, highlighting the sensitivity of the adult brain to social experience. Identifying the neural circuit mechanisms through which social experiences, such as isolation, modulate alcohol drinking will reveal how social context shapes vulnerability to alcohol misuse.

The basolateral amygdala (BLA) and medial prefrontal cortex (mPFC) are key brain regions in the distributed neural circuitry processing emotion, stress responses, and social information^17,18^, and their independent contributions to alcohol drinking are well established^19–22^. This led us to investigate whether BLA circuits adapt to social isolation in ways that influence alcohol use. Among these, BLA neurons projecting to the medial prefrontal cortex (BLA-mPFC) were of particular interest given their role in social behavior^23^ and negative valence^23–26^, pointing toward a potential involvement in aversive social contexts such as social isolation. Such contexts may shift the brain toward bottom-up processing, increasing the motivation for alcohol to alleviate negative emotional states. The BLA sends dense glutamatergic projections to the mPFC, targeting both excitatory and inhibitory neurons^27^. Activation of these BLA inputs predominantly suppresses mPFC activity^24,27,28^, suggesting a key modulatory role of this pathway in gating mPFC output. However, whether the BLA-mPFC circuit contributes to alcohol drinking, and how it adapts to aversive social experiences, remains unknown.

Using whole-cell patch-clamp electrophysiology, optogenetics, and cellular resolution calcium imaging, we tested the hypothesis that the BLA-mPFC circuit drives maladaptive alcohol drinking patterns in response to social isolation. We found that isolation increases alcohol drinking in males, while females show reduced alcohol intake. Yet in both sexes, BLA-mPFC excitability tracks alcohol drinking levels.

Given that social isolation selectively amplified drinking in males, but not females, we performed the remainder of our experiments primarily on males. Strikingly, the BLA-mPFC circuit robustly encodes and predicts alcohol drinking, and activation of this circuit increases alcohol drinking. During isolation, downstream mPFC neural representations shift, reducing responses to natural rewards such as sucrose while potentiating alcohol-related activity. Optogenetic stimulation of the BLA-mPFC circuit reproduced these shifts, mimicking an isolation-like cortical state, whereas inhibition of this pathway reduced alcohol consumption in isolated mice. Together, these findings reveal a BLA-mPFC circuit mechanism through which social isolation reconfigures prefrontal processing to promote alcohol use.

## RESULTS

### Social isolation increases alcohol drinking in males and decreases it in females

We first investigated how mice adapt their alcohol drinking in response to social isolation. We measured alcohol drinking in male and female mice using daily 1-hour two-bottle choice (15% alcohol vs. water) drinking sessions under group-housed conditions and across 11 days of social isolation (**Fig. 1a,b**).

**Figure 1.**
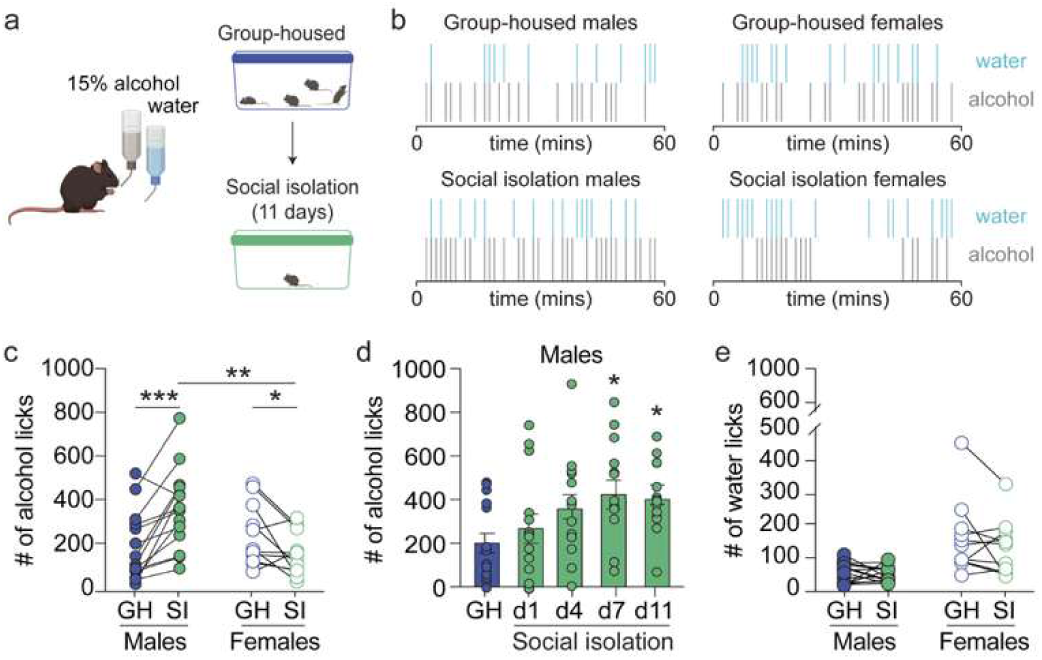
Social isolation increases alcohol drinking in males and decreases it in females. **a.** Schematic of the two-bottle choice paradigm in which mice had access to 15% alcohol and water before and during 11 days of social isolation created with BioRender. **b.** Representative alcohol licking raster plots from group-housed (top) and socially isolated (bottom) male (left) and female (right) mice during the drinking session. **c.** Social isolation produced opposite effects on alcohol licking in males and females (*N* = 14 male and 11 female mice, two-way ANOVA, interaction: F (1, 23) = 21.04, \*\*\**p* = 0.0001; Sidak’s multiple comparisons test), increasing alcohol licking in males while decreasing it in females. **d.** Time course of escalated alcohol drinking during social isolation (*N* = 14 male mice, one-way ANOVA, F (4, 65) = 2.83, \**p* = 0.03; post hoc Dunnett’s multiple comparisons test). **e.** Social isolation did not impact water licking, but females drink more water than males (*N* = 14 male and 11 female mice, two-way ANOVA, main effect of sex: F (1, 23) = 13.87, \*\**p* = 0.001). Some data points overlap due to similar values. Error bars indicate ±SEM.

A three-way ANOVA across sex, liquid, and housing condition revealed a significant interaction (F(1, 46) = 14.68; *p* < 0.001), indicating that social isolation altered alcohol and water drinking differently across sexes. Post hoc two-way analyses revealed that males exhibited a progressive escalation in alcohol intake across days of isolation, reaching a peak after approximately one week and remaining elevated thereafter (**Fig. 1c,d**). In contrast, isolation decreased alcohol drinking in females (**Fig. 1c**), revealing opposing behavioral adaptations to the same social experience. Water licking was higher in females overall but unaffected by housing condition (**Fig. 1e**).

Individual differences in alcohol intake were associated with social hierarchy in both sexes, and this relationship persisted during isolation (**Extended Data Fig. 1**). To verify that our behavioral measure accurately reflected intake, we compared lick counts to blood alcohol content and found a strong positive correlation (**Extended Data Fig. 1r**). Together, these findings demonstrate that social hierarchy contributes to variability in alcohol intake, and that social isolation selectively increases alcohol consumption in males, revealing a sex-dependent vulnerability to isolation-induced alcohol escalation.

### Social isolation increases BLA-mPFC neuronal excitability in males and decreases it in females

The BLA-mPFC circuit integrates affective and motivational signals to guide reward-related behaviors, suggesting it may be a key site where social isolation alters neural states that bias alcohol drinking. To test this, we examined how isolation impacts BLA-mPFC excitability. We performed whole-cell patch-clamp recordings from retrogradely labeled BLA-mPFC projection neurons throughout the basal nucleus of the BLA in group-housed and socially isolated male and female mice after 14 days (**Fig. 2a,b**).

**Figure 2.**
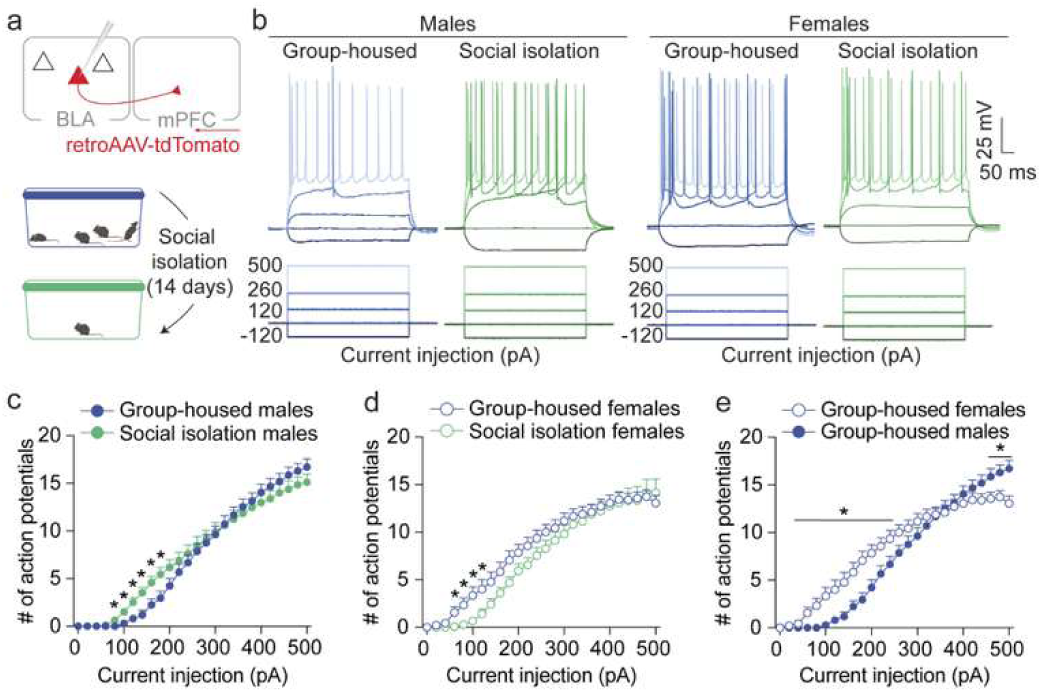
Social isolation increases BLA-mPFC neuronal excitability in males and decreases it in females. **a.** Viral strategy to label and measure excitability of BLA-mPFC neurons using *ex vivo* patch-clamp electrophysiology in group-housed and socially isolated mice created with BioRender https://BioRender.com/qsp4lsv. **b.** Representative action potential firing measured in BLA-mPFC neurons from a group-housed (blue) and socially isolated (green) male (left) and female (right) mice using current clamp recordings. **c.** Social isolation increases excitability of BLA-mPFC neurons in males (Group-housed: *n* = 21 neurons/*N* = 7 male mice; Social isolation: *n* = 30 neurons/*N* = 9 male mice, two-way ANOVA, interaction effect: F (25, 1225) = 4.3, \*\*\*\**p* < 0.0001; Sidak’s multiple comparisons post hoc, \**p* < 0.05). **d.** Social isolation decreases excitability of BLA-mPFC neurons in females (Group-housed: *n* = 18 neurons/*N* = 9 female mice; Social isolation: *n* = 15 neurons/*N* = 7 female mice, two-way ANOVA, interaction effect: F (25, 775) = 2.16, \*\*\**p* = 0.0009; Sidak’s multiple comparisons post hoc, \**p* < 0.05). **e.** Group-housed females exhibited higher baseline excitability of BLA-mPFC neurons compared to group-housed males (Group-housed females: *n* = 18 neurons/*N* = 9 female mice; Group-housed males: *n* = 21 neurons/*N* = 7 male mice, two-way ANOVA, interaction effect: F(25, 925) = 13.2, \*\*\*\**p* < 0.0001; Sidak’s post hoc, \**p* < 0.05). Error bars indicate ±SEM.

A three-way ANOVA across sex, current injection, and housing condition revealed a significant interaction (F(25, 2000) = 5.6; *p* < 0.0001), indicating that social isolation altered BLA-mPFC excitability differently across sexes. Post hoc two-way analyses showed that social isolation increased the intrinsic excitability of BLA-mPFC neurons in males, reflected by greater action potential firing across current injections (**Fig. 2c**) and elevated input resistance (**Table S1**), consistent with their greater alcohol drinking during isolation. Minimal effects of social isolation on excitability of non-specific BLA neurons were observed in males (**Extended Data Fig. 2**, **Table S1**), underscoring the selective sensitivity of the BLA-mPFC circuit to isolation.

In contrast to males, isolation decreased BLA-mPFC excitability in females (**Fig. 2d**), accompanied by reduced input resistance and a higher firing threshold (**Table S1**), consistent with their reduced alcohol drinking during isolation. Notably, females showed greater baseline excitability in BLA-mPFC compared to males (**Fig. 2e**), aligning with their trend in higher baseline alcohol intake.

Under group-housed conditions, BLA-mPFC neurons showed social rank-dependent differences in excitability (**Extended Data Fig. 2**; **Table S2**), paralleling rank-related variation in alcohol intake (**Extended Data Fig. 1**). These effects were not detected in non-specific BLA neurons (**Extended Data Fig. 3**; **Table S2**), suggesting that preexisting social dynamics may shape baseline BLA-mPFC activity.

Together, these findings demonstrate that the BLA-mPFC circuit consistently tracks alcohol drinking and represents a shared neural substrate of social isolation that adapts in opposite directions in males and females.

To further confirm that the BLA-mPFC circuit is functionally engaged during social isolation, we followed up on our previous finding that activating this pathway suppresses social interaction behavior^23^. Optogenetic inhibition of the BLA-mPFC circuit increased social interaction selectively during isolation (**Extended Data Fig. 4**), consistent with elevated circuit activity in vivo during this state and supporting its engagement in isolation-induced behavioral adaptations.

### BLA-mPFC activity robustly encodes and predicts alcohol drinking in males

Given the correlation between BLA-mPFC excitability and alcohol drinking patterns, we next asked how do BLA neurons respond to alcohol. In these and subsequent experiments, we focused on male mice, as they reliably escalate alcohol drinking during social isolation, offering a consistent behavioral phenotype for probing the underlying neural mechanisms.

Using cellular-resolution calcium imaging, we recorded the activity of both non-specific BLA neurons and projection-defined BLA-mPFC neurons in male mice as they consumed alcohol (**Fig. 3a**). We designed a drinking paradigm, cued two-bottle choice drinking (cued-2BC), that enabled trial-structured, cued availability of bottles (**Fig. 3b**). During the cued-2BC task, a cue light indicated the availability of alcohol and water bottles, allowing the mice approximately a minute of access before the bottles are retracted and an intertrial interval period begins. Note, we didn’t observe significant differences in alcohol drinking between GRIN implanted mice and controls, suggesting similar alcohol pharmacokinetics (**Extended Data Fig. 5a-c**).

**Figure 3.**
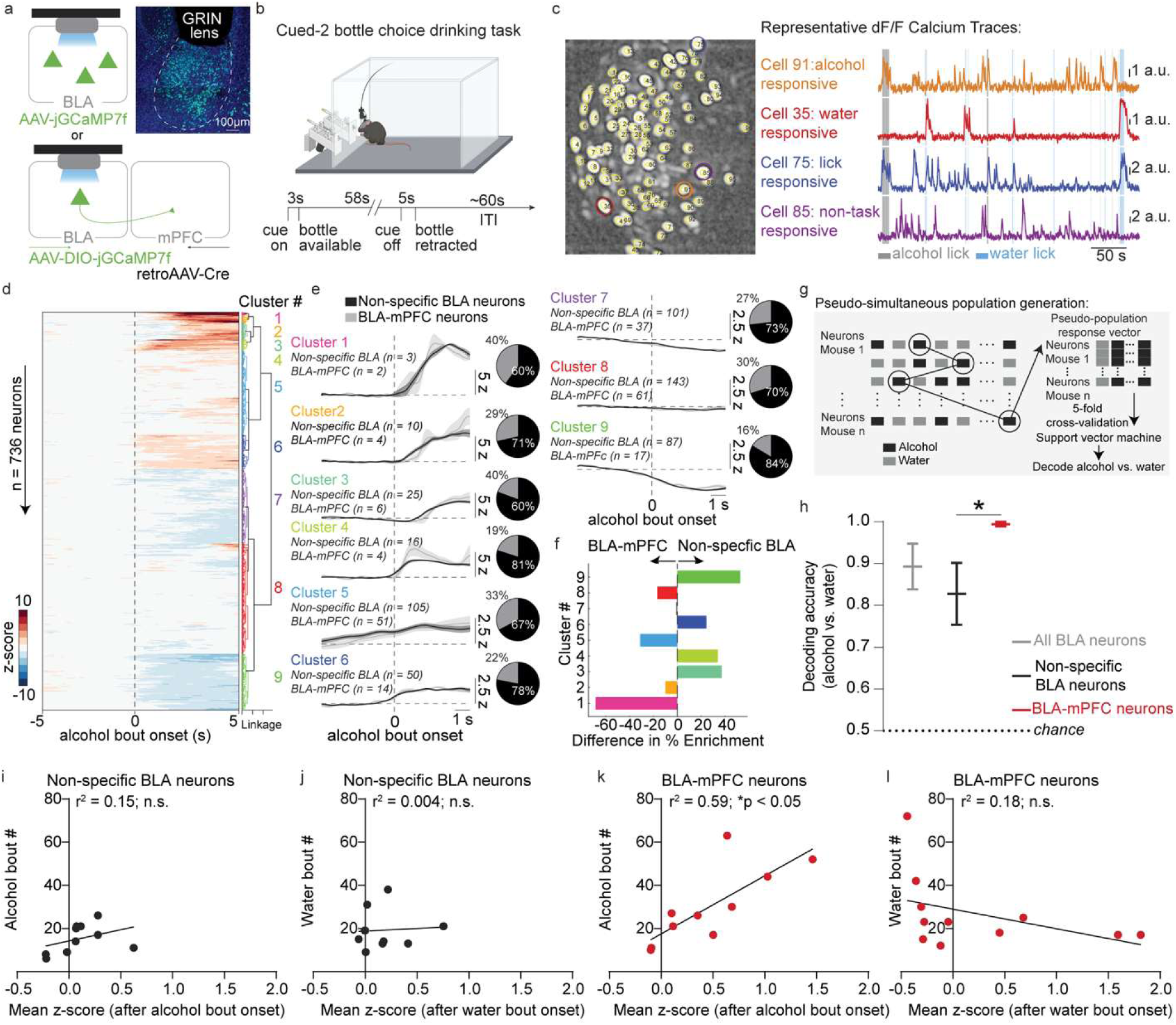
BLA-mPFC activity robustly encodes and predicts alcohol drinking in males. **a.** Viral strategy and GRIN lens implant for monitoring non-specific BLA and BLA-mPFC activity, replicated in all mice. **b.** Cued-two bottle choice (cued-2BC) drinking task, wherein a cue light signals availability of water and/or alcohol bottles for approximately a minute before bottles are retracted, created with BioRender https://BioRender.com/qsp4lsv. **c.** Representative cell contour map and calcium traces from BLA neurons during cued-2BC task. **d.** Functional activity clusters of non-specific BLA and BLA-mPFC alcohol responses (Non-specific BLA: *n* = 540 neurons/*N* = 10 male mice; BLA-mPFC projectors: *n* = 196 neurons/*N* = 11 male mice). **e.** Mean ±SEM z-score traces for functional clusters. Inset shows proportion of neurons per cluster for each population. **f.** Enrichment analysis showing distribution of BLA-mPFC and non-specific BLA neurons across functional clusters. **g.** Schematic of pseudo-simultaneous population sampling for decoding drinking from BLA population-level activity using a support vector machine. **h.** BLA-mPFC activity predicts alcohol drinking with greater accuracy than non-specific BLA neurons (Non-specific BLA: n = 540 neurons/N = 10; BLA-mPFC: n = 196 neurons/N = 11 male mice, Kruskal-Wallis test, H(2) = 8.06, \*\**p* = 0.009; Dunn’s multiple comparisons test, \**p* = 0.02). Mean ± SEM across 5 cross-validation folds. **i.** Non-specific BLA alcohol responses did not correlate with alcohol bouts (*N* = 10 male mice). **j.** Non-specific BLA water responses did not correlate with water bouts (*N* = 10 male mice). **k.** BLA-mPFC alcohol responses predict alcohol bouts (*N* = 11 male mice, two-sided Pearson correlation, *r*^2^ = 0.59, Bonferroni adjusted \**p* = 0.03). **l.** BLA-mPFC water responses did not correlate with water bouts (*N* = 11 male mice, two-sided Pearson correlation, *r*^2^ = 0.17, Bonferroni adjusted *p* = 0.76). Error bars indicate ±SEM.

Both non-specific BLA and BLA-mPFC neurons showed modest, transient increases in activity to the cue signaling alcohol and water availability (**Extended Data Fig. 5d**). Beyond this shared cue response, individual neurons exhibited diverse activity patterns, including selective responses to alcohol, water, general licking, or non-task-related events (**Fig. 3c,d**), indicating heterogeneous neural representations during drinking. Non-specific BLA and BLA-mPFC neurons excited to alcohol and water consisted of largely non-overlapping neurons (**Extended Data Fig. 5e,f**). Hierarchical clustering of responses to alcohol for all recorded BLA neurons resulted in several functional clusters (**Fig. 3d,e**). The distribution of BLA-mPFC neurons across functional clusters suggests that projection-defined BLA-mPFC neurons may represent a distinct functional subpopulation with specific response properties to alcohol, distinct from non-specific BLA neurons (**Fig. 3f**).

To assess whether BLA neurons encode alcohol-related information, we trained a support vector machine (SVM) classifier to distinguish activity during alcohol versus water consumption. Due to limited projection-defined neurons per mouse, we used a pseudo-simultaneous population sampling approach where drinking-associated time points were randomly sampled across all recorded neurons and mice to construct activity vectors (**Fig. 3g**). For five-fold cross-validation, each animal’s data was split into five parts, training and testing pseudo-populations were generated separately using the corresponding 80% and 20% of each animal’s data, and this was repeated across folds and iterations. While this method does not capture ensemble dynamics or neuron-neuron correlations, and restricts generalization across animals, it provides a useful approximation of population-level encoding by leveraging the full dataset to test whether distinct activity patterns are associated with each drinking condition.

We found that population activity in non-specific BLA neurons could distinguish alcohol from water consumption with ∼85% accuracy (**Fig. 3h**). Notably, decoding performance was even higher for projection-defined BLA-mPFC neurons (**Fig. 3h; Extended Data Fig. 6a,b**). These findings suggest that BLA-mPFC neurons contain a higher degree of alcohol-related information compared to non-specific BLA neurons, highlighting the potential importance of this circuit in alcohol drinking.

This led us to ask whether BLA-mPFC activity predicts alcohol intake. We correlated the number of alcohol bouts with the mean BLA-mPFC response to alcohol and found a significant positive relationship, such that greater BLA-mPFC activity predicted higher alcohol consumption (**Fig. 3k**). Average bout duration did not correlate with bout number (**Extended Data Fig. 6k**), indicating that BLA-mPFC activity tracks bout frequency rather than duration. No such relationship was observed for non-specific BLA neurons, water consumption, or cue responses (**Fig. 3i,j,l**; **Extended Data Fig. 6e-j**). Interestingly, when stratified by social rank, this relationship was significant only in subordinate mice (**Extended Data Fig. 6c,d**). Together, these findings indicate that BLA-mPFC activity encodes alcohol-related signals that predict drinking behavior.

### BLA-mPFC terminal stimulation increases alcohol drinking in males

To test whether BLA-mPFC activity causally modulates alcohol consumption, we used lick-triggered, closed-loop optogenetic activation of the BLA-mPFC circuit during two-bottle alcohol and water choice drinking (**Fig. 4a,b**). This design allowed circuit engagement to be precisely coupled to voluntary alcohol drinking. BLA-mPFC activation selectively increased alcohol intake, reflected by a significant rise in alcohol licks during laser stimulation that returned to baseline levels during recovery (**Fig. 4e**). In contrast, stimulation did not significantly alter licking for sucrose or water (**Fig. 4c,d**), indicating a specific enhancement of alcohol-related behavior.

**Figure 4.**
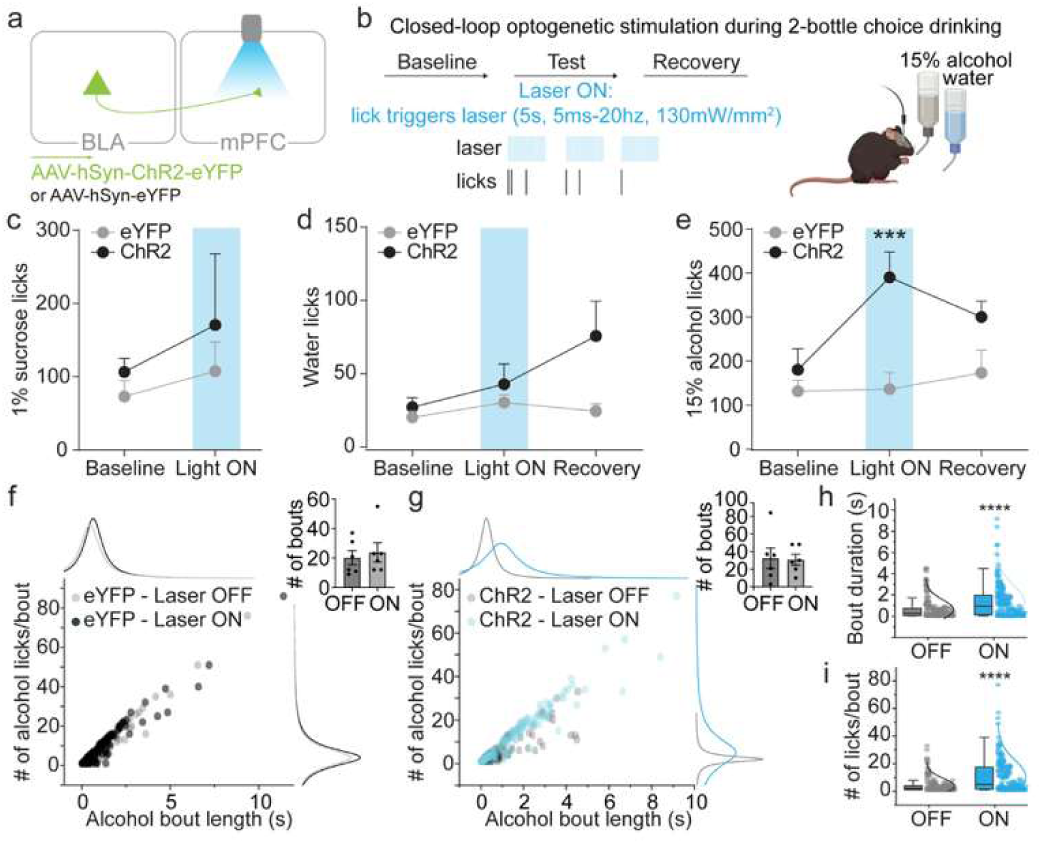
BLA-mPFC terminal stimulation increases alcohol drinking in males. **a.** Schematic of the viral strategy to express ChR2 or eYFP in BLA neurons projecting to the mPFC. **b.** To determine how BLA-mPFC circuits impact alcohol drinking under group-housed conditions, we used lick-triggered closed-loop optogenetics during two-bottle 15% alcohol and water choice drinking sessions, where licking at either bottle triggered 5 seconds of 20 Hz laser stimulation (130 mW/mm²), created with BioRender https://BioRender.com/qsp4lsv. **c, d.** BLA-mPFC terminal stimulation did not alter sucrose (c, *N* = 6 male mice/group) or water (d, *N* = 6 male mice/group) drinking. **e.** BLA-mPFC terminal stimulation increased alcohol drinking in ChR2-expressing males compared to eYFP controls (*N* = 6 male mice/group, two-way ANOVA, F (2, 20) = 4.7, \**p* = 0.02; Tukey’s multiple comparisons test). **f, g.** Distribution of alcohol bout lengths and number of licks per bout during non-stimulated (OFF) and BLA-mPFC stimulation (ON) conditions for eYFP control (f, *N* = 6 male mice) and ChR2 (g, *N* = 6 male mice) male mice. Insets show average number of bouts per condition. **h, i.** BLA-mPFC stimulation increased alcohol bout duration (h; *N* = 6 male mice, two-sided unpaired t-test, \*\*\*\**p* < 0.0001) and the number of alcohol licks per bout (i; *N* = 6 male mice, two-sided unpaired t-test, \*\*\*\**p* < 0.0001). Box plots display median (center line), 25th and 75th percentiles (box bounds), and 1.5× IQR (whiskers). Error bars indicate ±SEM.

Analysis of lick microstructure revealed that BLA-mPFC activation prolonged individual alcohol drinking bouts (**Fig. 4h**) and increased the number of licks per bout (**Fig. 4i**), without changing the total number of bouts (**Fig. 4g**). These effects were absent in eYFP controls (**Fig. 4f**), confirming that the behavioral changes were driven by circuit activation rather than nonspecific laser exposure. Together, these findings demonstrate that BLA-mPFC activation potentiates alcohol drinking by promoting longer, more sustained drinking bouts, suggesting this pathway may be a key driver of social isolation-linked increases in alcohol intake.

### Social isolation increases mPFC responses to alcohol while decreasing responses to sucrose in males

Despite the predictive power and causal role of BLA-mPFC activity in alcohol drinking, the precise mechanism by which this circuit influences alcohol drinking remained unclear. We hypothesized that the BLA may modulate mPFC representation of alcohol, which could alter motivation for alcohol. To investigate how the BLA impacts mPFC encoding of rewarding stimuli, we used a combined optogenetics and cellular resolution calcium imaging approach and measured mPFC dynamics during the cued-2BC drinking task before and following 14 days of social isolation in male mice **(Fig. 5a)**.

**Figure 5.**
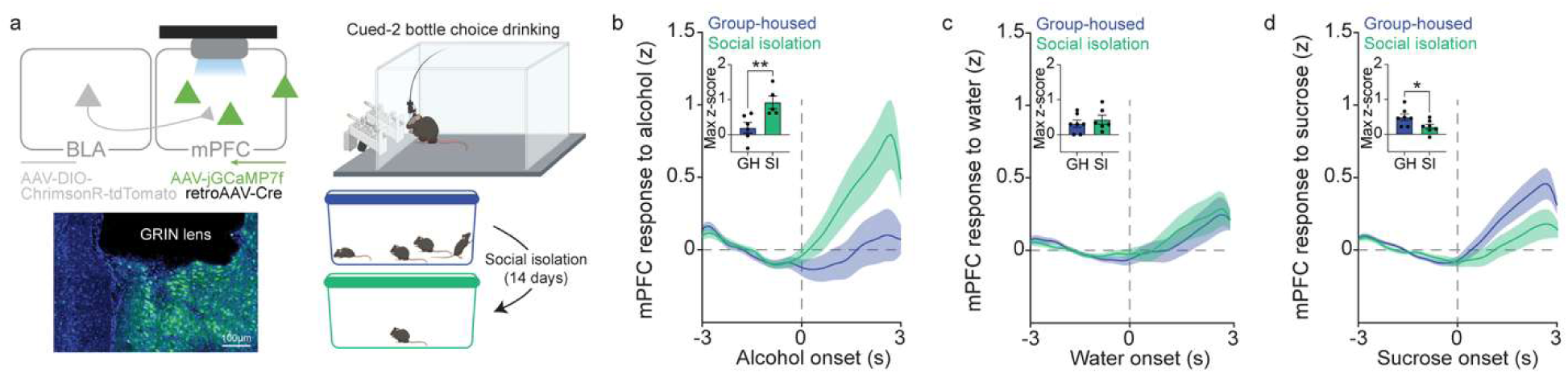
Social isolation increases mPFC responses to alcohol while decreasing responses to sucrose in males. **a.** Viral strategy and GRIN lens implant for simultaneous calcium imaging of mPFC neurons and optogenetic stimulation of BLA terminals in the mPFC, replicated in all mice. Recordings were conducted before and following 14 days of social isolation using the cued two-bottle choice paradigm, created in BioRender https://BioRender.com/qsp4lsv. **b.** Social isolation increased mPFC responses to alcohol (Group-housed: *n* = 240 neurons/*N* = 6 male mice; Social isolation: *n* = 281 neurons/*N* = 5 male mice, one-sided unpaired t-test, \*\**p* = 0.006). **c.** No detectable effect of social isolation was observed for mPFC responses to water (Group-housed: *n* = 240 neurons/*N* = 6 male mice; Social isolation: *n* = 281 neurons/*N* = 5 male mice). **d.** In contrast to alcohol, social isolation decreased mPFC responses to sucrose (Group-housed: *n* = 284 neurons/*N* = 7 male mice; Social isolation: *n* = 343 neurons/*N* = 7 male mice, one-sided unpaired t-test, \**p* = 0.01). Error bars indicate ±SEM.

We first examined how the mPFC responds to alcohol, water, and sucrose-related stimuli, and how these responses are altered by social isolation. mPFC responses to alcohol were minimal in group-housed mice but showed a marked increase during social isolation (**Fig. 5b**). Responses to water were unchanged (**Fig. 5c**). In contrast to alcohol, sucrose-evoked mPFC activity was reduced following isolation (**Fig. 5d**), consistent with a dampening of positive valence encoding. Importantly, population-level mPFC responses to alcohol and sucrose remained stable across 2-3 weeks of repeated imaging in group-housed mice (**Extended Data Fig. 7**), suggesting these changes reflect adaptations in neural representations by social isolation. Together, these findings demonstrate that social isolation reorganizes mPFC representations, diminishing natural reward while amplifying alcohol-related activity, potentially via modulation from the BLA, which regulates alcohol consumption.

### BLA-mPFC stimulation mimics social isolation-induced mPFC responses to alcohol and sucrose in males

To investigate how the BLA input impacts mPFC responses to alcohol, we stimulated BLA-mPFC terminals on 50% of trials in the cued-2BC drinking task (**Fig. 6a,b**) to allow comparison of mPFC dynamics with and without amplified BLA input.

**Figure 6.**
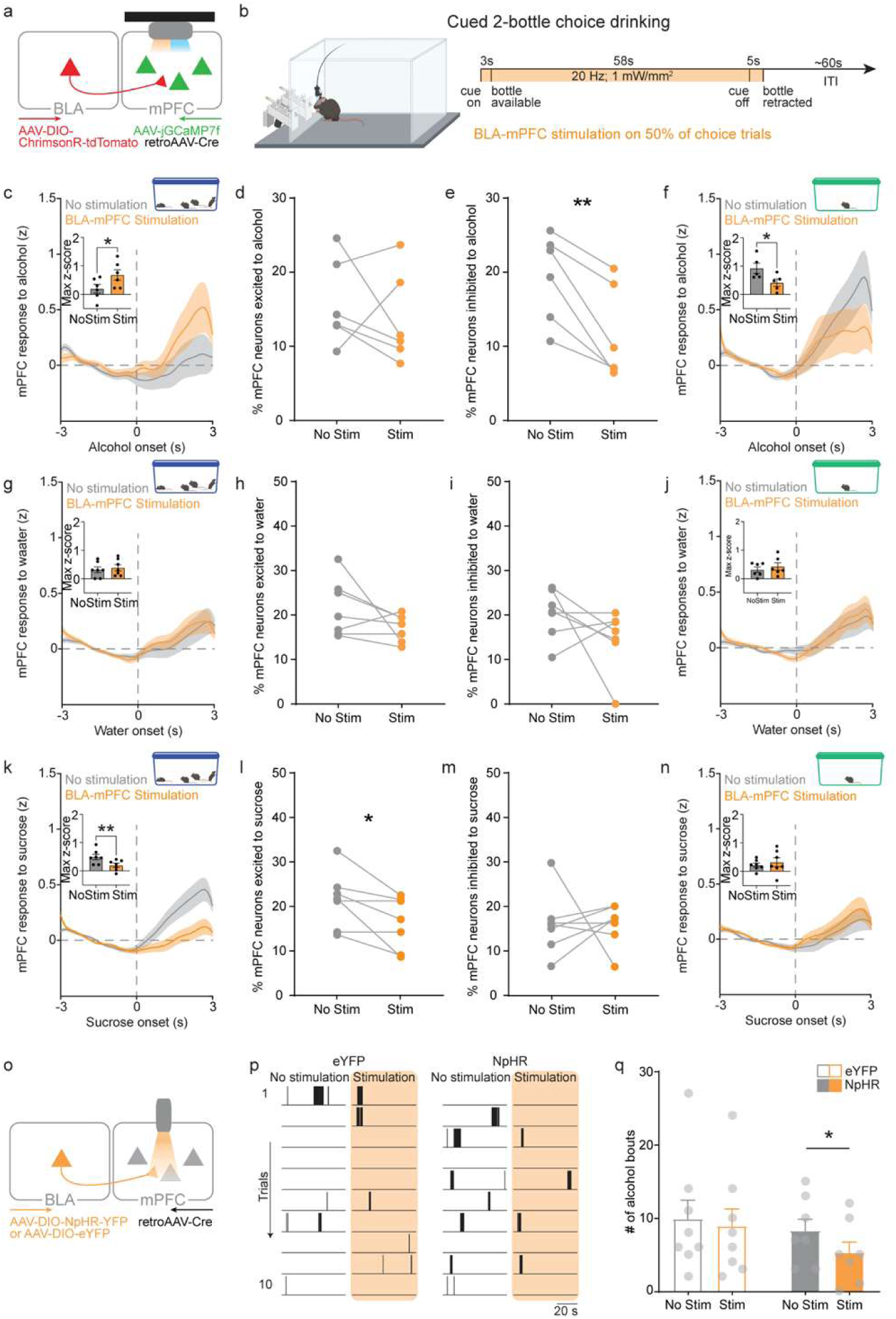
BLA-mPFC stimulation mimics social isolation-induced mPFC responses to alcohol and sucrose and inhibition decreases alcohol drinking during social isolation in males. **a.** Viral strategy and GRIN lens implant for mPFC imaging and BLA terminal stimulation. **b.** BLA-mPFC stimulation occurred during 50% of cued-2BC trials, before and during social isolation. **c.** BLA-mPFC stimulation increased mPFC alcohol responses in group-housed mice, mimicking social isolation (*n* = 240 neurons/*N* = 6 male mice, one-sided paired t-test, \**p* = 0.03). **d, e.** BLA-mPFC stimulation decreased inhibited mPFC alcohol responses (e; *N* = 6 male mice, one-sided paired t-test, t(5) = 4.6, \*\**p* = 0.005) without affecting excited responses (d); threshold of ±1.98. **f.** BLA-mPFC stimulation decreased mPFC alcohol responses during social isolation (*n* = 281 neurons/*N* = 5 male mice, one-sided paired t-test, \**p* = 0.02). **g-j.** BLA-mPFC stimulation had no effect on mPFC water responses in group-housed (g-i, *N* = 6) or social isolation (j, *N* = 5) male mice. **k.** BLA-mPFC stimulation decreased mPFC sucrose responses in group-housed mice, mimicking social isolation (*n* = 284 neurons/*N* = 7 male mice, one-sided paired t-test, \*\**p* = 0.008). **l, m.** BLA-mPFC stimulation decreased excited mPFC sucrose responses (l; *N* = 7 male mice, one-sided paired t-test, t(6) = 2.76, \**p* = 0.03) without affecting inhibited responses (m); threshold of ±1.98. **n.** Social isolation occluded BLA-mPFC stimulation effects on mPFC sucrose responses (*n* = 343 neurons/*N* = 7 male mice). **o.** Viral strategy to inhibit BLA-mPFC activity during 50% of cued-2BC trials during social isolation. **p.** Representative licking rasters for eYFP and NpHR mice. **q.** BLA-mPFC inhibition decreased alcohol bouts (*N* = 8 male mice/group, two-way ANOVA, main effect of stimulation: F(1,13) = 9.707, \*\**p* = 0.008; Sidak’s post hoc test, \**p* = 0.05). Error bars indicate ±SEM. Schematics created in BioRender https://BioRender.com/qsp4lsv.

We found that BLA-mPFC stimulation increased mPFC responses to alcohol (**Fig. 6c; Extended Data Fig. 8**), mimicking the effects of social isolation (**Fig. 5b**). Specifically, BLA-mPFC stimulation decreased the proportion of neurons inhibited in response to alcohol, without altering alcohol excited neurons (**Fig. 6d,e**). BLA-mPFC stimulation inhibited mPFC responses to alcohol during social isolation (**Fig. 6f; Extended Data Fig. 8**), pointing to neuroadaptations in this circuit by social isolation. Notably, no impact of BLA-mPFC stimulation on mPFC responses to water were observed (**Fig. 6g-j; Extended Data Fig. 8**).

Remarkably, BLA-mPFC stimulation was sufficient to also mimic the social isolation-induced decreases in mPFC responses to sucrose (**Fig. 6k**) and significantly decreased the proportion of neurons excited by sucrose, without altering neurons inhibited by sucrose (**Fig. 6l-m; Extended Data Fig. 8**). The effect of BLA-mPFC stimulation on mPFC responses to sucrose were occluded during social isolation (**Fig. 6n; Extended Data Fig. 8**), highlighting engagement of this mechanism by social isolation. Together, these data suggests that BLA-mPFC circuit stimulation recapitulates a social isolation-like state of mPFC encoding.

To further explore how BLA-mPFC activity influences cortical processing, we combined optogenetic BLA-mPFC terminal stimulation with calcium imaging of mPFC neurons during social interaction, tracking the same neurons across group-housed and social isolation conditions (**Extended Data Fig. 9**). Although not directly related to alcohol drinking, these experiments provided insight into how this pathway modulates mPFC ensemble dynamics across social states. In group-housed mice, BLA-mPFC stimulation reversed the polarity of mPFC responses to social contact, an effect absent after social isolation, indicating altered BLA-mPFC connectivity. Moreover, BLA-mPFC activation preferentially recruited neuronal ensembles resembling those engaged by social isolation, providing further support that this pathway can shift mPFC network states toward an isolation-like configuration.

### BLA-mPFC inhibition during social isolation reduces alcohol drinking in males

We then asked if the BLA-mPFC circuit causally regulates social isolation-associated drinking. To test this, we used optogenetics to inhibit the BLA-mPFC circuit on 50% of trials in the cued-2BC task during social isolation. We found that BLA-mPFC inhibition significantly decreased the number of alcohol bouts taken following social isolation, without altering water drinking (**Fig. 6o-q**), pointing toward a causal role of the BLA-mPFC circuit in social isolation-associated alcohol drinking. Taken together, these data suggest the BLA-mPFC circuit mimics a social isolation-like state of neural encoding and underlies behavioral adaptations in alcohol drinking.

## DISCUSSION

We defined a role for a subpopulation of BLA neurons projecting to the mPFC in driving alcohol consumption during social isolation. We found that individual variability in alcohol drinking is linked to social hierarchy in both sexes, and social isolation further increased alcohol drinking exclusively in male mice. We linked these behavioral phenomena to BLA-mPFC excitability, which reflects alcohol drinking patterns where both negative social contexts of social subordination and social isolation were associated with greater BLA-mPFC excitability in males. In line with these findings, we found that the BLA-mPFC circuit robustly encodes alcohol, as compared to non-specific BLA activity, and predicts alcohol drinking, where greater BLA-mPFC activity correlated with greater alcohol intake. Notably, mimicking BLA-mPFC activation was sufficient to increase alcohol drinking. In parallel to changes in the BLA-mPFC neurons, we found social isolation potentiated downstream mPFC responses to alcohol. Most notably, BLA-mPFC circuit stimulation mimicked the heightened mPFC alcohol responses observed during social isolation. Furthermore, inhibition of the BLA-mPFC circuit during social isolation reduced alcohol intake, suggesting a causal role of the BLA-mPFC circuit in alcohol drinking. Together, these findings underscore a key role of the BLA-mPFC circuit in mediating alcohol drinking in response to negative social contexts, providing insight into neural mechanisms by which social determinants shape vulnerability to an alcohol use disorder.

### Social influences on BLA-mPFC activity and alcohol drinking

While BLA-mPFC excitability emerged as a key circuit signature linking social context to alcohol consumption, behavioral responses to negative social contexts diverged by sex. Males escalated alcohol drinking whereas females decreased it, consistent with previous findings in rodents^29,30^ and mirroring human data^31,32^. This may reflect sex-specific processing of social isolation, as females show minimal behavioral or corticosterone changes after 14 days of isolation^33^. Despite these behavioral differences, BLA-mPFC excitability continued to scale with alcohol intake, showing higher excitability in isolated males and lower excitability in isolated females who drank more and less, respectively. Although the effects of isolation on intrinsic excitability appeared modest and were most evident at low current injection ranges, such near-threshold shifts can strongly influence synaptic integration and circuit recruitment. The bidirectional modulation of BLA-mPFC excitability across sexes may arise from differential regulation of calcium-activated or A-type potassium conductances that shape near-threshold firing. Notably, female mice also showed higher basal BLA-mPFC excitability than males, reflecting their greater baseline alcohol intake^34^. Together, these findings suggest that while the BLA-mPFC circuit tracks alcohol vulnerability in both sexes, its adaptation to social isolation is governed by sex-specific physiological mechanisms.

In addition to the challenges of social isolation, group living in social hierarchies presents its own social stressors that influence alcohol use. Across sexes, subordinates drank more alcohol than dominants, and baseline BLA-mPFC excitability was higher in lower-ranked males. Moreover, BLA-mPFC responses to alcohol were more strongly correlated with alcohol intake in lower ranked mice. This likely is motivated by the negative emotional states associated with social subordination rather than dominance or aggression^35–37^. In our study, social rank influenced alcohol but not water drinking, consistent with the idea that our limited-access paradigm engaged reward-driven rather than territorial motivation, which typically increases water drinking in dominant animals^38^. Together, these findings suggest that prior social experience and hierarchy history shape individual differences in alcohol-related behavior.

Notably, the heightened BLA-mPFC excitability observed in subordinate mice likely influences downstream mPFC activity, potentially contributing to the maintenance of social rank. Prior work shows that subordinates exhibit lower mPFC activity than dominants^39^, aligning with evidence that increased BLA activity predominantly inhibits mPFC neurons^24,27,28^. Together, these findings suggest that elevated BLA-mPFC excitability in subordinate mice may drive mPFC inhibition, thereby reinforcing lower social status.

Together, conceptually, our findings point to a shift toward bottom-up processing via the BLA-mPFC circuit during negative social contexts including social subordination and social isolation, which has previously been proposed to dominate during negative emotional states^40^. This consequently may lead to feedforward inhibition from BLA to mPFC to suppress top-down control, promoting maladaptive behaviors such as escalated alcohol intake. Consistent with this, reduced alcohol-related activity in mPFC neurons projecting to the periaqueductal grey has been linked to punishment-resistant, compulsive alcohol drinking phenotype^41^. In addition, activation of the BLA-mPFC circuit has been demonstrated to drive compulsive behavior^42^. Individual variability in alcohol consumption has also been linked to differences in behavioral flexibility^43^, which is regulated by the mPFC. Studies suggest that behavioral inflexibility not only predicts alcohol drinking but can be further impaired by alcohol exposure, sustaining maladaptive drinking behaviors^44^. This maladaptive alcohol drinking to cope with negative emotional states has been shown to be associated with an exacerbation of the subsequent transition to compulsive drinking with increased susceptibility to developing an AUD^45–47^.

### Impact of the BLA-mPFC circuit on mPFC encoding of natural rewards and alcohol

The mPFC is implicated in reward processing and valence encoding^18^. Our findings show that social isolation decreased mPFC responses to sucrose, consistent with negative affective states typically observed following social isolation^48^. Notably, BLA-mPFC stimulation was sufficient to drive decreased mPFC responses to sucrose. Specifically, BLA-mPFC stimulation led to a decrease in the percentage of mPFC cells activated by sucrose. This suggests that the BLA may suppress positive valence encoding in the mPFC, contributing to the affective deficits observed following isolation.

In contrast to natural rewards, we found that social isolation dramatically increased mPFC activity to alcohol. This increase is consistent with our finding that the BLA-mPFC circuit causally regulates alcohol drinking. In addition, we found that BLA-mPFC stimulation was sufficient to drive increased mPFC responses to alcohol. Notably, enhanced PFC response to alcohol associated cues is associated with relapse and craving in humans^49,50^. Reduced mPFC activity to natural rewards may contribute to the negative emotional states induced by social isolation, which can motive alcohol use as a coping mechanism. This, in turn, may potentiate mPFC responses to alcohol, reinforcing maladaptive alcohol drinking behavior. Dysregulated reward processing is a hallmark of substance use disorders as well as neuropsychiatric conditions such as depression and anxiety. By concurrently suppressing responses to natural rewards and amplifying responses to alcohol, the BLA-mPFC circuit may link social isolation to increased vulnerability to alcohol use.

The mPFC regulates social motivation, with its activity closely linked to social engagement^18^. Although not the primary focus of this study, we found that BLA-mPFC activity shapes mPFC responses to social contact. Isolation increased BLA-mPFC activity, and manipulating this pathway bidirectionally modulated social interaction, suggesting that elevated circuit activity during isolation contributes to reduced social motivation. At the network level, BLA-mPFC activation inverted responses of a subset of mPFC neurons to social contact, an effect abolished after isolation, likely reflecting diminished inhibitory gating, and recruited ensembles resembling those engaged during isolation. Together, these findings indicate that BLA-mPFC activity biases cortical states toward an isolation-like configuration, reducing social drive and reinforcing the behavioral effects of social isolation.

Together, our findings suggest that BLA input dynamically shapes mPFC activity and ensemble structure, biasing the network toward states that promote alcohol consumption. BLA-mPFC circuit activity not only scales with alcohol intake but also reorganizes mPFC population dynamics, favoring ensemble patterns that prioritize alcohol over natural rewards. By shifting the balance of mPFC encoding, the BLA-mPFC circuit may serve as a key node linking adverse social experience to altered motivation and maladaptive alcohol use.

## ACKNOWLEDGEMENTS

We would like to acknowledge the Scripps Research Animal Models Core for their measurements of blood alcohol content, the GENIE Project Team at HHMI Janelia Research Campus for developing the genetically encoded calcium indicator jGCaMP7f, and Dr. Nancy Padilla for helpful comments on the manuscript. Schematics created in BioRender. Wichmann, R. (2026) https://BioRender.com/qsp4lsv. This article is subject to HHMI’s Open Access to Publications policy. HHMI lab heads have previously granted a nonexclusive CC BY 4.0 license to the public and a sublicensable license to HHMI in their research articles. Pursuant to those licenses, the author-accepted manuscript of this article can be made freely available under a CC BY 4.0 license immediately upon publication.

## AUTHOR CONTRIBUTIONS

R.R.P and K.M.T. conceived of the project, designed and supervised the experiments. R.R.P. and K.K. performed stereotaxic surgeries. R.R.P., M.P., K.K, R.P., A.H., R.C., A.P., and X.Y. performed behavioral experiments. R.R.P., B.J., and J.W. performed patch-clamp experiments. R.R.P., K.K, R.P., A.H., performed calcium-imaging experiments. R.R.P., M.P., K.K, R.P., A.H., H.L., and C.C. analyzed the data. F.H.T. and M.K.B. performed computational analysis. H.L., R.C., and A.H. performed SLEAP automated pose tracking analysis. R.R.P., M.P., K.K, R.P., and H.L. performed and analyzed optogenetic experiments. R.R.P., M.P., K.K, R.P., and R.C. performed histological verifications. R.R.P., H.L., C.R.L., C.J. and L.K. provided MATLAB scripts and advice for data analysis. C.C., R.L.M., C.J., J.D., K.B., A.L., T.D.P. and R.W. made additional significant intellectual contributions. R.R.P. graphed data and made figures. R.R.P and K.M.T. wrote the paper. All authors contributed to editing the manuscript.

## METHODS

### Animals and housing

Adult male and female C57BL/6J mice (≥10 weeks) from Jackson Laboratory were housed 3-4/cage on a 12-h reverse light/dark cycle with ad libitum food and water. All experiments were conducted during the dark phase. All experimental procedures were carried out in accordance with NIH guidelines and approval of the Salk Institutional Animal Care and Use Committee and Northwestern University Institutional Animal Care and Use Committee

### Stereotaxic surgeries

All surgeries were conducted under aseptic conditions. Mice were anesthetized with isoflurane (4-5% induction, 1-2% maintenance), placed in a stereotaxic frame (David Kopf Instruments), and given subcutaneous lidocaine (0.5%) at the shaved, betadine/alcohol-cleaned incision site. Eye lubricant was applied and body temperature maintained with a heating pad. A midline incision exposed the skull and a craniotomy was performed. All mice received subcutaneous Ringer’s solution (1 mL), buprenorphine (1 mg/kg), and meloxicam (5 mg/kg) intraoperatively, recovered on a heating pad, and given ≥14 days before behavioral testing.

All stereotaxic coordinates were measured relative to bregma and the top of the skull. Viral vectors were injected through glass pipettes pulled to a 100-200 µm tip diameter, attached to either a microsyringe pump (Hamilton Microlitre 701, Hamilton Co.) or Nanoject III programmable nanoliter injector (Drummond Scientific). Pipettes were slowly lowered to the target site and virus delivered at 1.0 nL/s. After 2 min, the pipette was raised 0.02 mm and left in place for 8 min to allow diffusion before slow withdrawal.

For non-specific BLA calcium imaging, 200 nL of AAV1-Syn-jGCaMP7f-WPRE (Addgene) was injected into the BLA (AP: −1.35, ML: +3.4, DV: −5.0 mm) and a 0.6 mm diameter × 7.3 mm GRIN lens with integrated baseplate (Inscopix) was implanted above the BLA (DV: −4.8 mm).

For projection-specific BLA-mPFC calcium imaging, AAV1-Syn-FLEX-jGCaMP7f-WPRE (200 nL) was injected into the BLA (AP: −1.35, ML: +3.4, DV: −5.0 mm), retrograde AAV2-hSyn-Cre-P2A-tdTomato (200 nL) into the mPFC (AP: +1.9, ML: +0.4, DV: −1.9 mm), and a 0.6 mm × 7.3 mm GRIN lens with integrated baseplate was implanted above the ipsilateral BLA.

For simultaneous mPFC calcium imaging and BLA terminal stimulation, a mixture of AAV1-Syn-jGCaMP7f-WPRE and retrograde AAV2-hSyn-Cre-P2A-tdTomato (100 nL each) was injected into the mPFC (AP: +1.9, ML: +0.4, DV: −1.9 mm), AAV8-hSyn-Flex-ChrimsonR-tdTomato (200 nL; Addgene) into the ipsilateral BLA (AP: −1.35, ML: +3.4, DV: −5.0 mm), and a 0.5 mm × 4 mm GRIN lens with integrated baseplate implanted above the ipsilateral mPFC (DV: −1.7 mm). No tissue was aspirated. All lens implants were secured with adhesive cement (C&B Metabond, Parkell) followed by black cranioplastic cement (Ortho-Jet, Lang) and the incision closed with nylon sutures.

To label BLA-mPFC neurons for ex vivo electrophysiology, retrograde AAV2-CAG-tdTomato (200 nL) was injected into the right mPFC (AP: +1.9, ML: +0.4, DV: −1.9 mm).

For BLA-mPFC optogenetic excitation during the resident-intruder task, retrograde AAV2-hSyn-Cre-P2A-tdTomato (200 nL) was injected unilaterally into the mPFC (AP: +1.9, ML: +0.4, DV: −1.9 mm) and AAV8-hSyn-Flex-ChrimsonR-tdTomato or AAV8-Flex-tdTomato (200 nL) into the ipsilateral BLA (AP: −1.35, ML: +3.4, DV: −5.0 mm). A 300 µm optic fiber (0.37 NA, 1.25 mm ferrule) was implanted 0.30 mm above the mPFC injection site.

For BLA-mPFC optogenetic excitation during two-bottle choice drinking, AAV5-hSyn-ChR2-eYFP or AAV5-hSyn-eYFP (200 nL) was injected unilaterally into the BLA (AP: −1.35, ML: +3.4, DV: −5.0 mm) and a 300 µm optic fiber (0.37 NA, 1.25 mm ferrule) implanted 0.30 mm above the mPFC injection site.

For BLA-mPFC optogenetic inhibition during the resident-intruder task and cued two-bottle choice drinking, retrograde AAV2-hSyn-Cre-P2A-tdTomato (200 nL) was injected bilaterally into the mPFC and AAV5-EF1-DIO-NpHR-eYFP or AAV5-EF1-DIO-eYFP (200 nL) into the BLA (same coordinates as above). A single 300 µm optic fiber (0.37 NA, 1.25 mm ferrule) was implanted 0.30 mm above the midline mPFC (AP: +1.9, ML: 0.0, DV: −1.6 mm).

All fiber implants were secured with adhesive cement followed by black cranioplastic cement, and the incision closed with nylon sutures once dry.

### Behavioral assays

All behavioral testing occurred after ≥2 weeks post-surgery recovery. Mice were handled 15 min/day for 3-5 days prior to testing to reduce experimenter-induced stress.

### Tube dominance test

Social rank was assessed using the tube dominance test. Mice were habituated to a short Plexiglas tube (7.5 cm × 4.5 cm ID) in their home cage for 1-3 days, then trained to traverse a longer tube (30 cm × 3.2 cm ID) over ≥3 days. Mice then competed in 3-5 daily round-robin trials against each cagemate, released simultaneously from opposite ends. The retreating mouse was designated the loser. Relative social rank was defined as the proportion of wins across 3-4 days of testing and considered stable when consistent for ≥4 consecutive days.

### Two-bottle choice drinking

Alcohol drinking was assessed using daily 1-hour two-bottle choice (2BC) sessions in operant chambers (Med Associates) controlled by Med-PC IV 4.2 software, equipped with white noise and lickometer-wired bottles. A sucrose fade initiated drinking: mice first received water and 10% sucrose until >500 licks on consecutive days, then progressed through 10% sucrose + 15% alcohol (days 1-3), 2% sucrose + 15% alcohol (days 4-6), and 15% alcohol alone (day 7 onward). Following 5-6 baseline days, mice were socially isolated for 11-14 days with continued daily 2BC. Alcohol licks were averaged over the last 3 baseline days and across 11 isolation days. Mice with <50 alcohol licks were excluded from calcium imaging analyses.

### Calcium imaging during cued two-bottle choice drinking

To examine neural activity during alcohol drinking, we developed a trial-structured cued-2BC task. Operant chambers were equipped with white noise, retractable lickometer-wired bottles, and a cue light above each bottle. A 3-s cue preceded bottle availability (58-s access); the cue extinguished 5 s before retraction. Intertrial intervals were 53-73 s (randomized). Each session comprised 30 trials: 20 choice (both bottles available), 5 forced water, and 5 forced alcohol/sucrose.

The 2BC sucrose fade procedure was used to initiate drinking, with the following modifications. Following sucrose-only 2BC, mice were trained on cued-2BC with sucrose for 5 days before the sucrose fade. After baseline alcohol 2BC, calcium imaging recordings were obtained using using Inscopix Data Acquisition Software during alcohol cued-2BC sessions. Mice were mounted with a miniscope or dummy miniscope for all sessions.

### Optogenetic stimulation and calcium imaging during cued two-bottle choice drinking

The cued-2BC procedure was used with the following modifications. After baseline, an additional cued-2BC session was given in which stimulation (20 Hz, 590–650 nm, 1 mW/mm²) was delivered on 50% of choice trials in randomized order, from cue onset until bottle retraction. Mice were then isolated and given alcohol 2BC sessions. After 14 days of isolation, mice received reminder cued-2BC sessions without stimulation followed by stimulation sessions for both alcohol and sucrose. Calcium imaging was obtained during all stimulation sessions. Mice were mounted with a miniscope or dummy miniscope and received one session per day.

### Optogenetic inhibition during cued two-bottle choice drinking

The cued-2BC procedure was used as described above. Due to low drinking in this cohort, mice received 2 weeks of drinking-in-the-dark (DID) prior to the inhibition session: daily 2-h home cage access to 15% alcohol and water for 4 days, 4-h access on day 5, and 2 days off, repeated for 2 weeks. An inhibition cued-2BC session was then conducted following baseline and isolation cued-2BC, in which continuous photoinhibition (589 nm, 15 mW at fiber tip) was delivered on 50% of randomized choice trials from cue onset until bottle retraction. Mice were tethered for all sessions.

### Optogenetic stimulation during two-bottle choice drinking

Following baseline 2BC, a test session was given in which lick onset at either spout triggered 5 s of blue light stimulation (20 Hz, 479 nm, 9 mW at fiber tip), followed by a no-stimulation recovery session. Mice then received a baseline session of 1% sucrose and water 2BC followed by a stimulation test session. Mice were tethered for all sessions.

### Optogenetic stimulation during resident-juvenile intruder task

Mice were tested individually in their home cage with a 3-min habituation period, followed by a 6-min interaction with a novel juvenile during which stimulation was delivered for the first 3 min and withheld for the final 3 min. The task was repeated after 14 days of social isolation with a different juvenile. Stimulation parameters: ChrimsonR, 20 Hz, 589 nm, 15 mW at fiber tip; NpHR, continuous, 589 nm, 15 mW at fiber tip. Behavior was recorded using Noldus EthoVision XT 14, and social interaction during the 3-min stimulation epoch was quantified using SLEAP automated pose tracking (described below).

### Optogenetic stimulation and calcium imaging during resident-juvenile intruder task

Mice were mounted with miniscopes (nVoke, Inscopix) and given a 5-min habituation in their home cage, followed by a 5-min interaction with a novel juvenile during which stimulation (20 Hz, 590-650 nm, 1 mW/mm²) or no stimulation was delivered, counterbalanced across two consecutive days. A novel juvenile was used for each session. This procedure was repeated after 14 days of social isolation. Social interaction during the 5-min interaction period was quantified using SLEAP automated pose tracking (described below).

### Lick microstructure analysis

Lick bouts were defined as ≥2 consecutive licks within 1 s, with bout onset at the first lick. Licks per bout and total bouts per session were calculated using custom code in Matlab 2025b. For calcium imaging analyses, only bouts separated by ≥3 s were included.

### Calcium imaging analysis

#### Calcium imaging data acquisition and calcium signal extraction

Calcium imaging data were collected using nVista and nVoke systems (Inscopix). Miniscope recordings were triggered via TTL signal for the duration of each behavioral session, with the miniscope connected to an active commutator (Inscopix).

Raw videos were processed in IDAS (Inscopix): spatially downsampled 4x, temporally downsampled to 10 Hz, cropped, and motion-corrected to the first frame. Videos were exported as TIFF stacks and converted to 8-bit in Fiji. Fluorescence traces were extracted using constrained non-negative matrix factorization algorithm optimized for micro-endoscopic imaging (CNMF-E)^51^ without non-negative constraints on temporal components to preserve negative transients associated with decreases in firing. All neurons were visually curated by experimenters blinded to experimental condition; neurons with abnormal morphology or calcium traces were excluded.

### Population-level mean response calculation

Neuronal responses to drinking and social interaction bouts were calculated by z-score normalizing each neuron’s GCaMP7f signal to a 3-or 5-s pre-bout baseline: Z = (F(t) − F_m)/SD, where F_m is the mean ΔF/F₀ during the baseline period. Z-scored traces were averaged across a matched number of bouts per condition per neuron, then across all neurons to obtain the population mean response. Responsive neurons were identified by Wilcoxon signed-rank test or a mean Z-score of ±1.98 from bout onset to 3 s.

For resident-juvenile intruder recordings, neurons were co-registered across sessions using CellReg, developed by Sheintuch *et al.*^52^, which aligns CNMF-E spatial footprints to a reference session via rotational and translational shifts and identifies cell pairs using a Bayesian method based on centroid distance and spatial correlation.

### Agglomerative hierarchical clustering

Agglomerative hierarchical clustering (Ward linkage, Euclidean distance) was applied to each neuron’s mean z-score trace. The linkage threshold was set to the highest value yielding at least one non-responsive cluster. Neurons within each cluster were averaged to generate peri-event time traces. For co-registered neurons, pre-and during-isolation mean z-score responses were horizontally concatenated prior to clustering, and cluster averages were computed separately for each epoch.

### Pseudo-simultaneous population generation and decoding for alcohol drinking

In each experimental session and for every individual, drinking bouts lasting more than 1s were identified, with each bout being distinct and separated by a minimum interval of 2 seconds from another. Bouts were categorized based on whether the individual was consuming alcohol (or sucrose) or water, thereby defining our two conditions.

Following the identification, we performed a k-fold cross-validation procedure. This procedure was executed ensuring that each fold contained at least one bout from each class. By ensuring this representation in every fold, we aimed to guarantee an unbiased and comprehensive evaluation of our machine learning models.

Subsequently, we generated pseudo-simultaneous training and testing populations from the respective datasets of all subjects^53,54^. In this context, a pseudo-simultaneous population refers to an aggregated or amalgamated set of neuronal activity data gathered from multiple animals. The purpose of this construction is to simulate a larger, collective neuronal population, enabling more robust computational or statistical analysis.

The process of creating this pseudo-simultaneous population involved several steps. First, for each subject, we pooled neuronal activity data corresponding to the identified bouts of drinking behavior. Next, we sampled a behavioral label (0 or 1) and pseudo-randomly sampled a single timepoint from each subject corresponding to that behavioral label. The neural activity of all animals can then be stacked creating a single vector of shape 1 x N, where N is the total number of neurons across all animals. This was then repeated T times (T=10,000 unless otherwise noted) to create a pseudo-simultaneous population of shape T x N. We then followed the same procedure to create a testing population using the testing data.

The resultant training and testing pseudo-simultaneous populations were then used to train and evaluate our linear Support Vector Machine classifiers. The entire procedure was repeated for ’k’ iterations, as defined in the k-fold cross-validation scheme. The training data was balanced by subsampling the majority-class before training the model to avoid introducing bias.

Creating the pseudo-simultaneous population for the social interaction data was identical to the method outlined above. However, instead of separating the data into drinking bouts, it was separated into bouts of interaction, and bouts of noninteraction. Noninteraction bouts that were longer than 20 seconds were divided into multiple smaller bouts while still separating each bout by at least 2 seconds.

### Enrichment Analysis

Percent enrichment was calculated by comparing observed to expected cell counts per cluster, where expected counts were determined by proportional distribution based on population size. Positive and negative values indicate enrichment and depletion, respectively. Differences in percent enrichment between populations were plotted.

### SLEAP automated pose tracking analysis

Behavioral videos were acquired using a Basler GenICam camera at 25 fps with Noldus EthoVision XT 14, with cameras positioned at a fixed height above the cage for all recordings. Animal poses were estimated using SLEAP^55^, with a 6-point skeleton (nose, left/right ears, skull base, haunch, tail base) labeled on 798 frames per model to train bottom-up models for ChrimsonR and NpHR experiments. All frames were visually confirmed for correct identity assignment. Social interaction was defined by a resident head-to-intruder body distance of ≤60 pixels and angle of ≤135°, validated against manually scored data and consistent with unsupervised clustering results (**Extended Data Fig. 4**).

For calcium imaging experiments, social interaction was identified using SLEAP-predicted body point coordinates to calculate pairwise features between the resident and intruder: head-to-head distance and angle, resident head-to-intruder tail base distance and angle, intruder head-to-resident tail base distance and angle, tail base-to-tail base distance, and each animal’s velocity. Uniform Manifold Approximation and Projection (UMAP) was applied to these features to obtain unsupervised behavioral clusters, which were manually annotated as social or non-social based on visualization of representative videos (**Videos S1** and **S2**).

### Ex vivo whole-cell patch-clamp electrophysiology

A separate cohort was injected to label BLA-mPFC projection neurons for patch-clamp recordings. Following 4 weeks of recovery, mice underwent tube dominance testing to establish stable social ranks, then were either isolated for 14-16 days or remained group-housed. Viral injection sites in the mPFC were visually confirmed before recordings.

Mice were anesthetized with sodium pentobarbital (200 mg/kg, i.p.), decapitated, and brains extracted in oxygenated (95% O2/5% CO2) high-sucrose cutting solution (in mM: 87 NaCl, 2.5 KCl, 1.3 NaH2PO4, 7 MgCl2, 25 NaHCO3, 75 sucrose, 5 ascorbate, 0.5 CaCl2; ∼300-320 mOsm, pH 7.3). Coronal slices (300 µm) containing the mPFC and BLA were collected on a vibrating microtome and incubated in oxygenated ACSF (in mM: 126 NaCl, 2.5 KCl, 1.25 NaH2PO4, 1 MgCl2, 2.4 CaCl2, 26 NaHCO3, 10 glucose; ∼300 mOsm, pH 7.3) at 31°C for 1 h. Slices were visualized under an upright microscope (Scientifica) with IR-DIC optics.

Whole-cell patch-clamp recordings were obtained from tdTomato-expressing BLA-mPFC neurons and adjacent non-expressing neighbors using 3-5 MΩ glass pipettes filled with internal solution (in mM: 125 potassium gluconate, 20 HEPES, 10 NaCl, 3 MgATP, 8 biocytin; ∼290 mOsm, pH 7.3). Signals were acquired, amplified, and digitized at 10 kHz (MultiClamp 700B, Digidata1440b, pClamp10; Molecular Devices). Slices were continuously perfused with oxygenated ACSF at 31 ± 1°C throughout recordings.

Intrinsic excitability was assessed by holding neurons at −70 mV and applying 500-ms current injections from −120 to +500 pA in 20-pA increments. Action potential number was plotted against current injection to generate excitability curves, with firing threshold defined as the minimum current eliciting an action potential. Cell capacitance and resting membrane potential were recorded directly; input resistance was estimated from the steady-state response to a −120 pA hyperpolarizing step. Data were analyzed using Clampfit v.11 (Molecular Devices) and GraphPad Prism 10. Recording pipette location within the slice was imaged through a 4× /0.10 NA objective.

### Blood alcohol concentration

Blood alcohol concentration was assessed to determine if measurements of alcohol licking correlated with alcohol consumption. Following experimental testing, terminal trunk blood was collected from mice immediately after completion of a two-bottle choice drinking session. Blood was collected in an Eppendorf tube and immediately centrifuged to separate serum. Blood alcohol content was measured on an Agilent 7820A GC coupled to a 7697A (headspace-flame-ionization) through the Scripps Research Animal Models Core.

### Histology

Mice were deeply anesthetized with sodium pentobarbital (200 mg/kg, i.p.) and transcardially perfused with Ringer’s solution followed by cold 4% PFA/PBS. For viral injection verification, brains were extracted and post-fixed in 4% PFA/PBS for 24 h at 4°C. For optic fiber and GRIN lens verification, whole heads were submerged in 4% PFA/PBS for 24 h at 4°C, after which implants were removed and brains extracted. Brains were cryoprotected in 30% sucrose/PBS at 4°C, sectioned coronally at 50 µm, mounted, and coverslipped with DAPI-containing mounting medium. Sections were imaged at 2× (Keyence BZ-X Wide Image Viewer 1.3.1.1). Mice with incorrect viral expression or misplaced implants were excluded. Injection sites and implant placements are annotated in **Extended Data Fig. 10**.

### Statistics

All analyses were performed in GraphPad Prism and MATLAB. No statistical methods were used to pre-determine sample sizes, but our sample sizes are similar to those reported in previous publications. Two-group comparisons used paired or unpaired one-or two-tailed t-tests; three or more groups used one-, two-, or three-way ANOVA with repeated measures as appropriate. Parametric or non-parametric tests were selected based on normality and homogeneity of variance assumptions. Pearson’s correlation was used for two-variable associations with multiple comparisons correction. Fisher’s exact test and chi-square assessed distributional differences. Significance threshold: p < 0.05. Data are presented as mean ± SEM unless otherwise stated. Mice were randomly assigned to groups and optogenetic stimulation trials were randomized. Experimenters were blinded during analysis but not during data collection.

## DATA AVAILABILITY

All data supporting the findings of this study are available in the main text, supplementary materials, and the source data files provided with this paper. Unprocessed data have been deposited in the DANDI Archive under accession code 001454 (https://dandiarchive.org/dandiset/001454).

## CODE AVAILABILITY

All custom code used in this study has been deposited in the DANDI Archive under accession code 001454 (https://dandiarchive.org/dandiset/001454).

## EXTENDED DATA

**Extended Data Figure 1.**
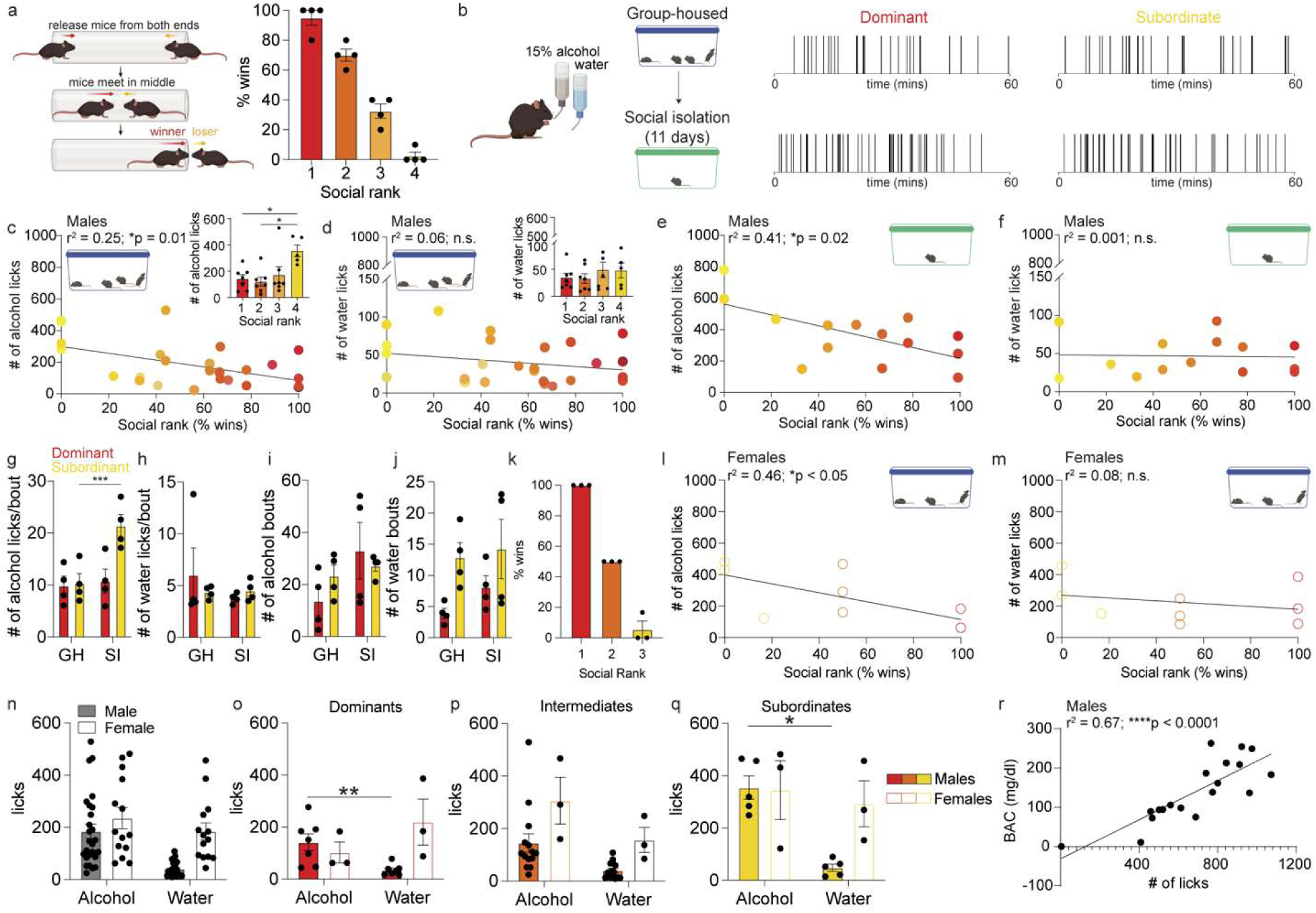
Alcohol, but not water, drinking varies with social rank in both sexes, with females showing greater water consumption. **a.** Percent wins per rank in male mice. **b.** Representative alcohol licking rasters from a dominant and subordinate male mouse during two-bottle choice drinking before and during social isolation. **c.** Rank correlates with alcohol licks (*N* = 26 male mice, Pearson correlation, *r*^2^ = 0.25, Bonferroni adjusted \**p* = 0.01). Inset, Rank 4 mice drink more alcohol (*N* = 26 male mice, one-way ANOVA, F(3, 22) = 4.68, \**p* = 0.01; Sidak’s post hoc, \**p* < 0.05). **d.** No correlation between social rank and water licks (*N* = 26 male mice, Pearson correlation, *r*^2^ = 0.06, *p* = 0.43). Inset, No rank differences in water drinking. **e, f.** Rank correlates with alcohol licks during isolation (e; *N* = 14 male mice, Pearson correlation, *r*^2^ = 0.41, Bonferroni adjusted \**p* < 0.05), but not water (f; *N* = 14 male mice, Pearson correlation, *r*^2^ = 0.002, Bonferroni adjusted *p* = 1). **g, h.** Subordinate males increase alcohol licks per bout during isolation (g; *N* = 4 male mice/group, two-way ANOVA, interaction: F (1, 6) = 20.84, \*\*\**p* < 0.003, main effect of isolation: F(1, 6) = 28.84, \*\**p* < 0.01; Sidak’s post hoc, \*\*\**p* = 0.001 compared to group-housed), with no effect on water licks per bout (h). **i, j.** Isolation increases alcohol bout number in dominant males (i; *N* = 4 male mice/group, two-way ANOVA, main effect of isolation: F (1, 6) = 10.23, \**p* = 0.01), with no effect on water bouts (j). **k.** Percent wins per rank in female mice (*N* = 9 mice). **l,m.** Rank correlates with alcohol licks (*N* = 9 female mice, one-tailed Pearson correlation, based on male mice, *r*^2^ = 0.46, Bonferroni adjusted \**p* = 0.04), but not water licks (m, Pearson correlation, *r*^2^ = 0.08, Bonferroni adjusted *p* = 0.45). **n.** Males show higher alcohol and lower water licking than females (*N* = 26 male and 14 female mice/group, two-way ANOVA, main effect of sex: F(1, 38) = 10.77, \*\**p* = 0.002; main effect of liquid: F(1, 38) = 17.29, \*\*\**p* = 0.0002). **o-q.** Alcohol and water licking by sex in dominants (o; *N* = 7 male and 3 female mice, two-way ANOVA, interaction: F(1, 8) = 7.05, \**p* = 0.05, Sidak’s post hoc, \*\**p* < 0.01), intermediates (p; *N* = 14 male and 3 female mice, 2 two-way ANOVA, main effect of sex: F(1, 15) = 7.99, \**p* = 0.01, main effect of liquid: F(1, 15) = 10.1, \*\**p* = 0.006), and subordinates (q; *N* = 5 male and 3 female mice, two-way ANOVA, interaction: F(1, 6) = 10.96, \**p* = 0.01, Sidak’s post hoc, \**p* < 0.05). **r.** Alcohol licks (15% alcohol + 10% sucrose) correlates with blood alcohol concentrations, (*N* = 19 male mice, Pearson’s correlation, *r*^2^ = 0.67, \*\*\*\**p* < 0.0001). Some data points overlap. Error bars indicate ±SEM. Schematics created in BioRender https://BioRender.com/qsp4lsv.

**Extended Data Figure 2.**
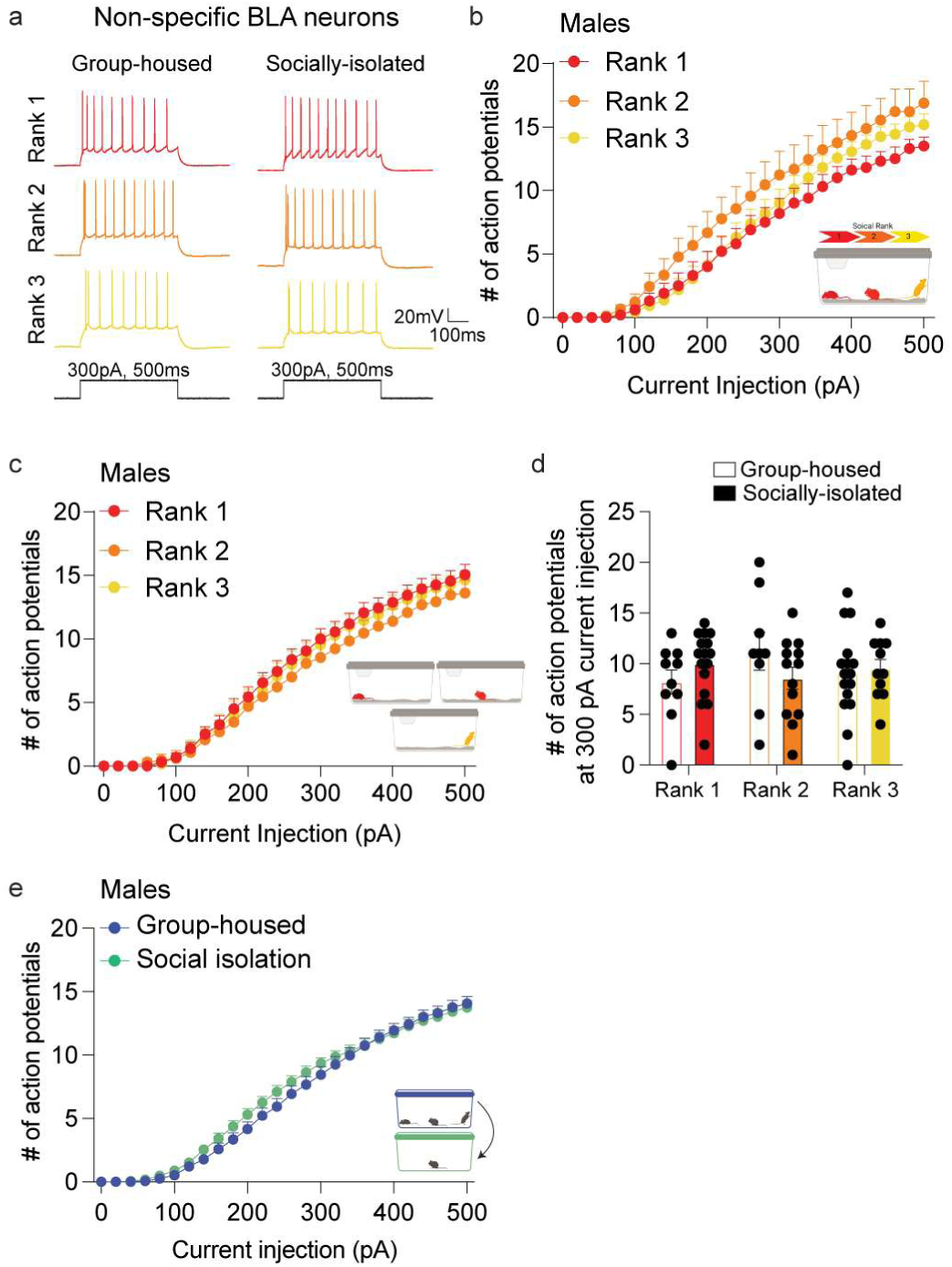
Non-specific BLA neurons show increased excitability following social isolation, with no effect of social rank in males. **a.** Representative action potential firing measured in non-specific BLA neurons from group-housed and 14 days socially isolation male mice for each social rank. **b, c.** No detectable social rank-dependent differences were observed in non-specific BLA neurons from group-housed (b; Rank 1: *n* = 10 cells from *N* = 3 male mice; Rank 2: *n* = 9 cells from *N* = 3 male mice; Rank 3: *n* = 16 cells from *N* = 3 male mice) or socially isolated male mice (c; Rank 1: *n* = 16 cells from *N* = 3 male mice; Rank 2: *n* = 13 cells from *N* = 3 male mice; Rank 3: *n* = 11 cells from *N* = 3 male mice). **d.** Social isolation did not alter the number of action potentials elicited by a 300 pA current injection across each social rank in males. **e.** Social isolation increases excitability of non-specific BLA neurons in males (Group-housed: *n* = 50 neurons from *N =* 11 male mice; Social isolation: *n* = 54 neurons from *N =* 12 male mice; two-way ANOVA, interaction effect: F (25, 2550) = 1.58, \**p* = 0.03). Error bars indicate ±SEM. Schematics created in BioRender https://BioRender.com/qsp4lsv.

**Extended Data Figure 3.**
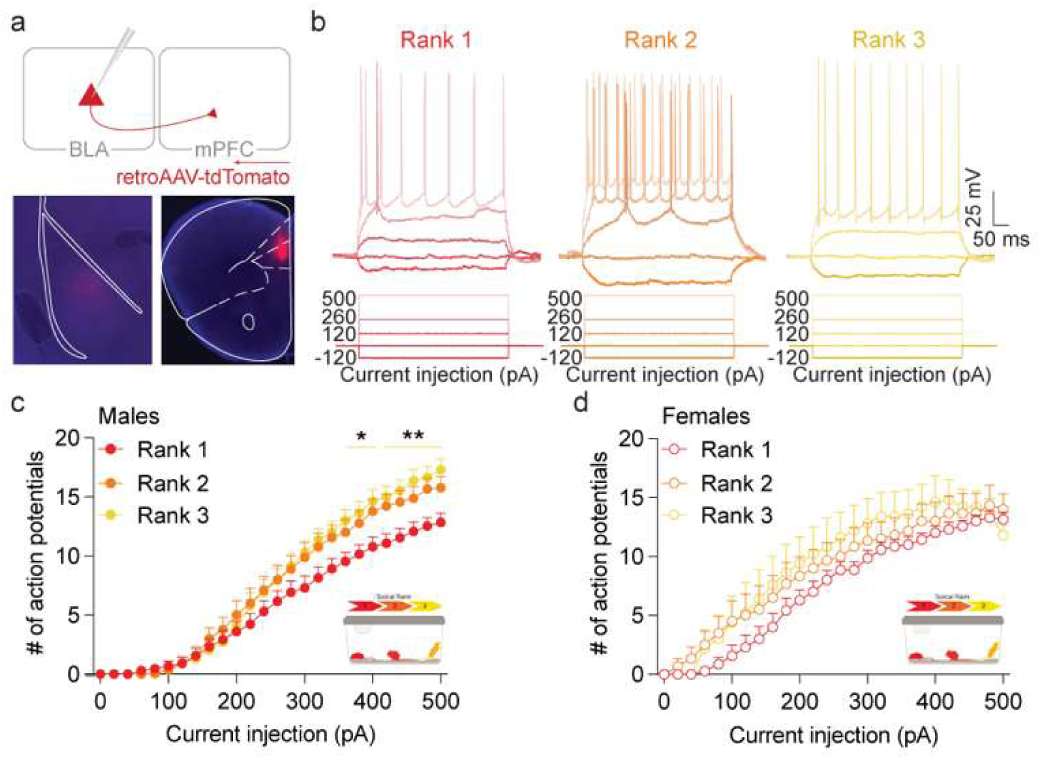
BLA-mPFC neuronal excitability varies with social rank in a sex-specific manner. **a.** Viral strategy to label and measure excitability of BLA-mPFC neurons using whole-cell patch-clamp electrophysiology across social rank, with representative post-patching images showing targeted neuronal labeling, replicated in all mice. **b.** Representative action potential firing measured in BLA-mPFC neurons from Dominant (Rank 1, red), Intermediate (Rank 2, orange), and Subordinate (Rank 3, yellow) male mice using current-clamp recordings. **c.** Dominant male mice showed reduced basal BLA-mPFC excitability compared to Intermediate and Subordinate male mice (Rank 1: *n* = 13 neurons from *N* = 4 male mice; Rank 2: *n* = 9neurons from *N* = 4 male mice; Rank 3: *n* = 14 neurons from *N* = 5 male mice, two-way ANOVA, interaction effect: F (50, 825) = 4.25, \*\*\*\**p* < 0.0001; Tukey’s post hoc, \**p* < 0.05, \*\**p* < 0.01, compared to Rank 1). **d.** No detectable social rank-dependent differences were observed in BLA-mPFC neurons from group-housed female mice (Rank 1: *n* = 7 neurons from *N* = 3 female mice; Rank 2: *n* = 6 neurons from *N* = 3 female mice; Rank 3: *n* = 5 neurons from *N* = 3 female mice). Error bars indicate ±SEM. Schematics created in BioRender https://BioRender.com/qsp4lsv.

**Extended Data Figure 4.**
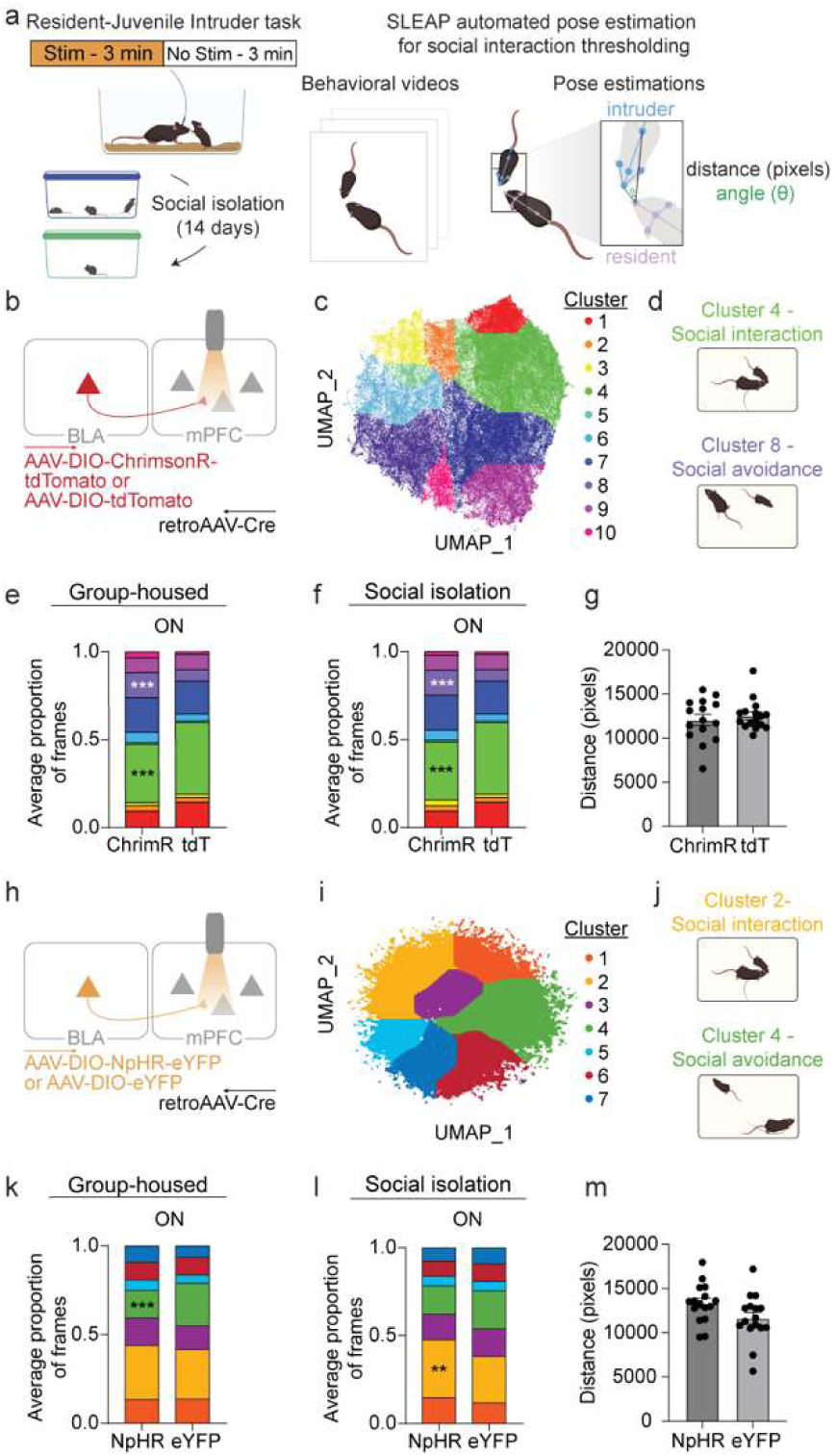
BLA-mPFC manipulation bidirectionally regulates social interaction during isolation as identified by behavioral clustering in males. **a.** Schematic of optogenetic stimulation epochs during the resident-juvenile intruder task before and after isolation. SLEAP pose estimation identified social interaction frames. **b.** Viral strategy for BLA terminal activation in mPFC. **c.** Unsupervised UMAP clustering of behavioral frames from ChrimsonR and tdTomato male mice. **d.** Schematic of frames from social interaction and social avoidance clusters. **e,f.** BLA-mPFC stimulation decreased social interaction in ChrimsonR vs. tdTomato males under group-housed (e; two-way ANOVA, interaction: F(9,140) = 4.94, \*\*\*\**p* < 0.0001; Sidak’s post hoc, \*\*\**p* < 0.001) and social isolation (f; interaction: F(9,140) = 4.99, \*\*\*\**p* < 0.0001; Sidak’s post hoc, \*\*\**p* < 0.001) conditions (*N* = 8 male mice/group). **g.** Total distance traveled did not differ between ChrimsonR and tdTomato mice (ChrimsonR: *N* = 15; tdTomato: *N* = 16 male mice). **h.** Viral strategy for BLA terminal inhibition in mPFC. **i.** Unsupervised UMAP clustering of behavioral frames from NpHR and eYFP male mice. **j.** Schematic of frames from social interaction and social avoidance clusters. **k.** BLA-mPFC inhibition decreased social avoidance in NpHR vs. eYFP males under group-housed (k; two-way ANOVA, interaction: F(6,98) = 4.02, \*\**p* = 0.001; Sidak’s post hoc, \*\*\**p* < 0.001) and increased social interaction under social isolation (l; two-way ANOVA, interaction: F(6,97) = 3.57, \*\**p* = 0.003; Sidak’s post hoc, \*\**p* < 0.01) conditions (*N* = 8 male mice/group). **m.** Distance traveled did not differ between NpHR and eYFP mice (*N* = 16 male mice/group). Error bars indicate ±SEM. Schematics created in BioRender https://BioRender.com/qsp4lsv.

**Extended Data Figure 5.**
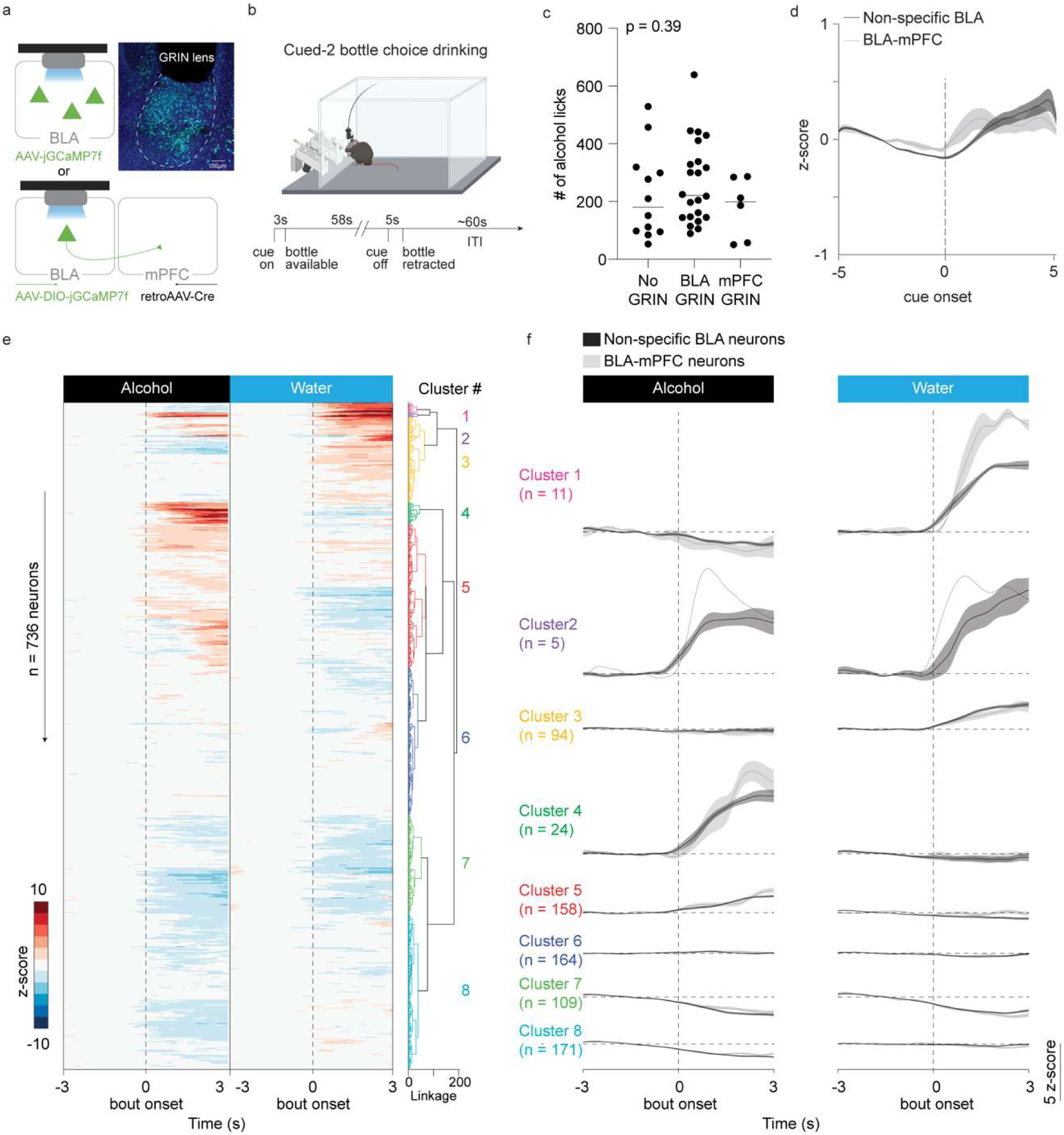
Non-specific BLA and BLA-mPFC neurons excited to alcohol and water consisted of largely non-overlapping ensembles in males. **a.** Viral strategy and GRIN lens implant for monitoring non-specific BLA and projection-specific BLA-mPFC neuronal activity, replicated in all mice. **b.** To record in vivo BLA dynamics during drinking, we designed a cued-two-bottle choice drinking task to enable conditioned availability of bottles and free-choice drinking, wherein a cue light signals availability of water and/or alcohol bottles for approximately a minute after which the bottles are retracted, created in BioRender.com. **c.** GRIN lens implants do not alter baseline alcohol licks (No GRIN: *N* = 12 male mice; BLA GRIN: *N* = 22 male mice; mPFC GRIN: *N* = 6 male mice, one-way ANOVA, F (2, 37) = 0.47, *p* = 0.62). **d.** Mean calcium responses of non-specific BLA and BLA-mPFC neuronal populations to the cue signaling alcohol or water availability. Both populations exhibited modest, transient increases in activity (<0.5 z-score) following cue onset (Non-specific BLA: *n* = 540 neurons from *N* = 10 male mice; BLA-mPFC projectors: *n* = 196 neurons from *N* = 11 male mice). **e.** Functional activity clusters of combined non-specific BLA and BLA-mPFC neuronal responses to alcohol and water (Non-specific BLA: *n* = 540 neurons from *N* = 10 male mice; BLA-mPFC projectors: *n* = 196 neurons from *N* = 11 male mice). **f.** Mean responses of each functional cluster stratified by non-specific BLA and BLA-mPFC neuronal populations. Error bars indicate ±SEM.

**Extended Data Figure 6.**
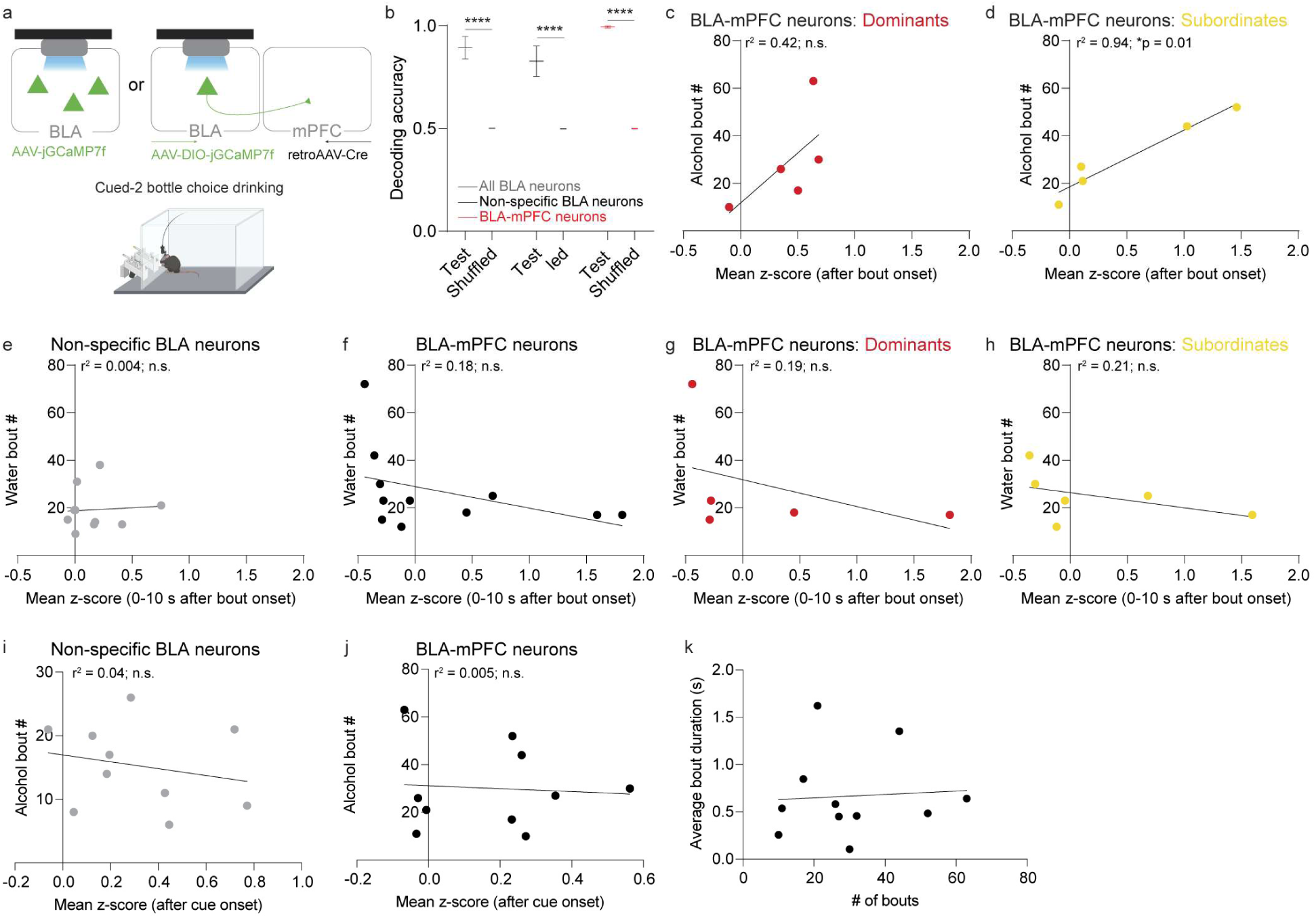
BLA-mPFC activity predicts alcohol, but not water, drinking in a rank-dependent manner in males. **a.** Strategy for monitoring non-specific BLA and BLA-mPFC neuronal activity during the cued-2BC task (BioRender https://BioRender.com/qsp4lsv). **b.** Population-level activity of non-specific BLA, BLA-mPFC, and all BLA neurons decodes alcohol vs. water drinking above chance (shuffled data), using a support vector machine (Non-specific BLA: *n* = 540 neurons/*N* = 10 male mice; BLA-mPFC: *n* = 196 neurons/*N* = 11 male mice; two-way ANOVA, main effect: F(1,24) = 174.2, \*\*\*\**p* < 0.0001; Sidak’s post hoc, \*\*\*\**p* < 0.0001). **c,d.** BLA-mPFC alcohol responses correlate with alcohol bout number in lower-ranked (Ranks 3-4) males (d; Pearson correlation, *r*^2^ = 0.94, Bonferroni adjusted \**p* = 0.01) but not higher-ranked (Ranks 1-2) males (c; *r*^2^ = 0.42, Bonferroni adjusted *p* = 1). **e, f.** Non-specific BLA (e; Pearson correlation, *r*^2^ = 0.004, *p* = 0.84) and BLA-mPFC (f; Pearson correlation, *r*^2^ = 0.17, Bonferroni adjusted *p* = 1) activity was not correlated with water drinking. **g, h.** No correlation was observed between BLA-mPFC activity and water drinking in Dominants (g; Pearson correlation, *r*^2^ = 0.19, Bonferroni adjusted *p* = 1) or Subordinates (h; Pearson correlation, *r*^2^ = 0.21, Bonferroni adjusted *p* = 1) male mice. **i, j.** Scatter plots showing non-specific BLA neurons (i, *r*^2^ = 0.04, *p* = 0.53) and BLA-mPFC neurons (j, *r*^2^ = 0.005, *p* = 0.84) cue-evoked responses did not correlate with alcohol bouts. **k.** No correlation was observed between the average bout duration and the number of bouts (*N* = 11 male mice, Pearson correlation, *r*^2^ = 0.0, *p* = 0.84). Error bars indicate ±SEM.

**Extended Data Figure 7.**
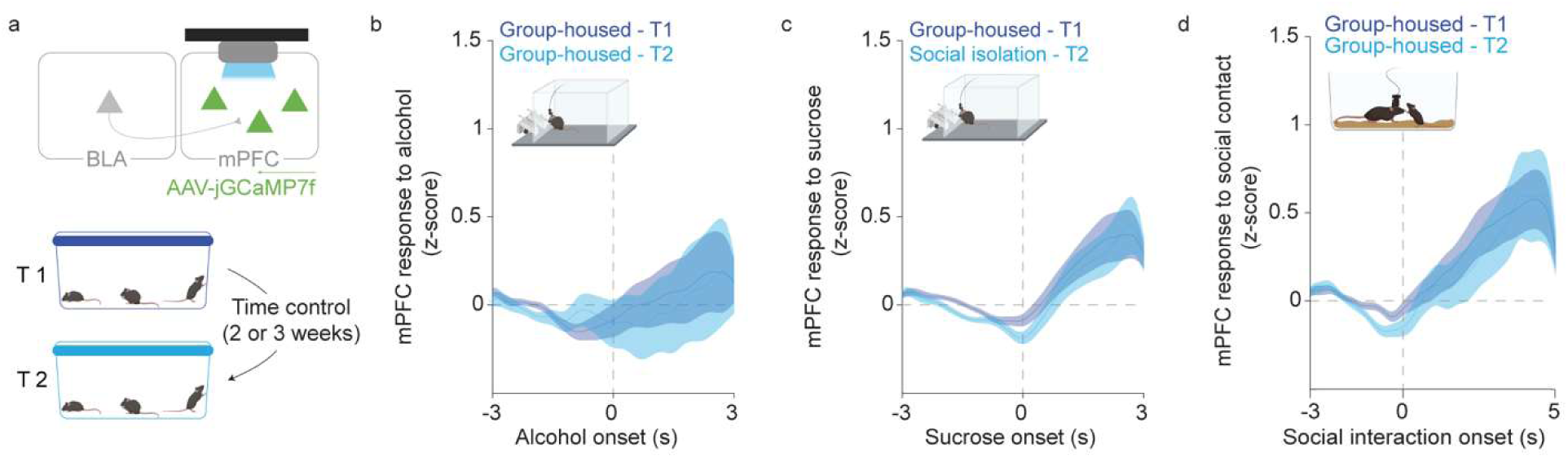
mPFC responses to alcohol, sucrose, and social contact are stable across time in males. **a.** Viral strategy and GRIN lens implant for monitoring of mPFC activity. Calcium imaging was performed in group-housed male mice at two distinct timepoints separated by 2-3 weeks. **b.** No effect of time on mean mPFC responses to alcohol drinking in the cued 2-bottle choice drinking task (T1: *n* = 275 neurons and T2: *n* = 275 neurons from *N* = 4 male mice). **c.** No effect of time on mean mPFC responses to sucrose (T1: *n* = 165 neurons and T2: *n* = 195 neurons from *N* = 5 male mice). **d.** No effect of time on mean mPFC responses to social interaction during the resident-intruder task (T1: *n* = 237 neurons and T2: *n* = 168 neurons from *N* = 6 male mice). Error bars indicate ±SEM. Schematics created in BioRender https://BioRender.com/qsp4lsv.

**Extended Data Figure 8.**
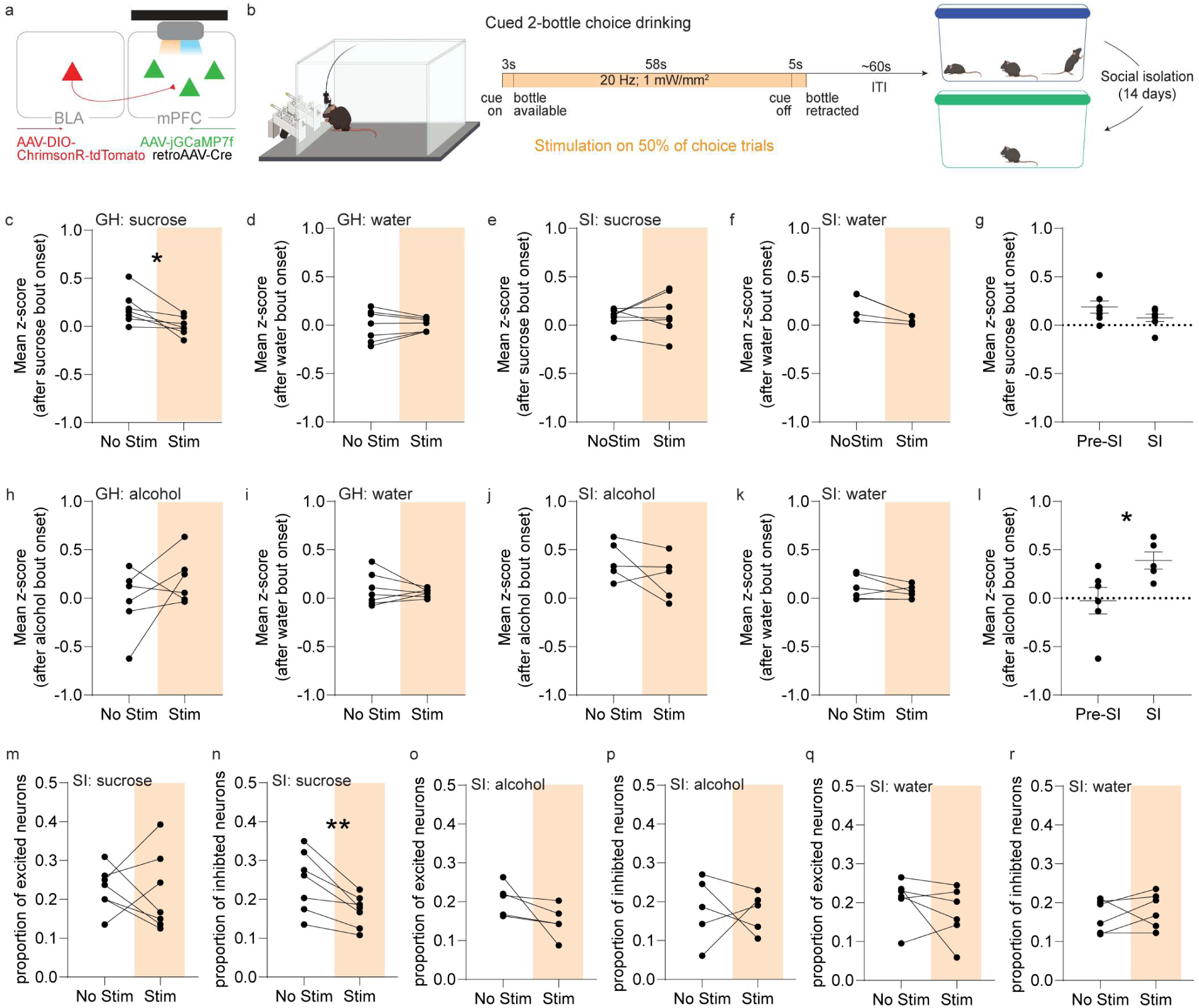
BLA-mPFC stimulation alters mPFC responses and the proportion of responsive neurons to sucrose, alcohol, and water during group housing and social isolation in males. **a.** Viral strategy and GRIN lens implant for simultaneous monitoring of mPFC activity and stimulation of BLA terminals in the mPFC. **b.** To measure the impact of BLA stimulation on mPFC dynamics during drinking, we used the cued-two-bottle choice drinking task, wherein BLA-mPFC stimulation occurred during 50% of trials, which was repeated before and after 14 days of social isolation, created in BioRender https://BioRender.com/qsp4lsv. **c, d.** BLA-mPFC stimulation significantly decreased the mean mPFC response to sucrose (c; paired t-test, \**p* = 0.02) with no effect on mPFC responses to water (d) under group-housed conditions. **e, f.** BLA-mPFC stimulation did not alter mPFC responses to sucrose (e) or water (f) during social isolation. **g.** Mean mPFC responses to sucrose before and during social isolation was not significantly different. **h, i.** BLA-mPFC stimulation did not alter mPFC responses to alcohol (h) or water (i) pre-social isolation. **j, k.** BLA-mPFC stimulation did not alter mPFC responses to alcohol (j) or water (k) during social isolation. **l.** Social isolation significantly increased the mean mPFC responses to alcohol (unpaired t-test, \**p* = 0.03). **m, n.** BLA-mPFC stimulation significantly decreased the proportion of mPFC neurons inhibited to sucrose (n; paired t-test, \*\**p* = 0.005) with no effect on mPFC neurons excited to sucrose (m), based on a mean Z-score response ±1.98, during social isolation. **o, p.** BLA-mPFC stimulation did not alter the proportion of mPFC neurons responsive to alcohol during social isolation. **q, r.** BLA-mPFC stimulation did not alter the proportion of mPFC neurons responsive to water during social isolation. Error bars indicate ±SEM.

**Extended Data Figure 9.**
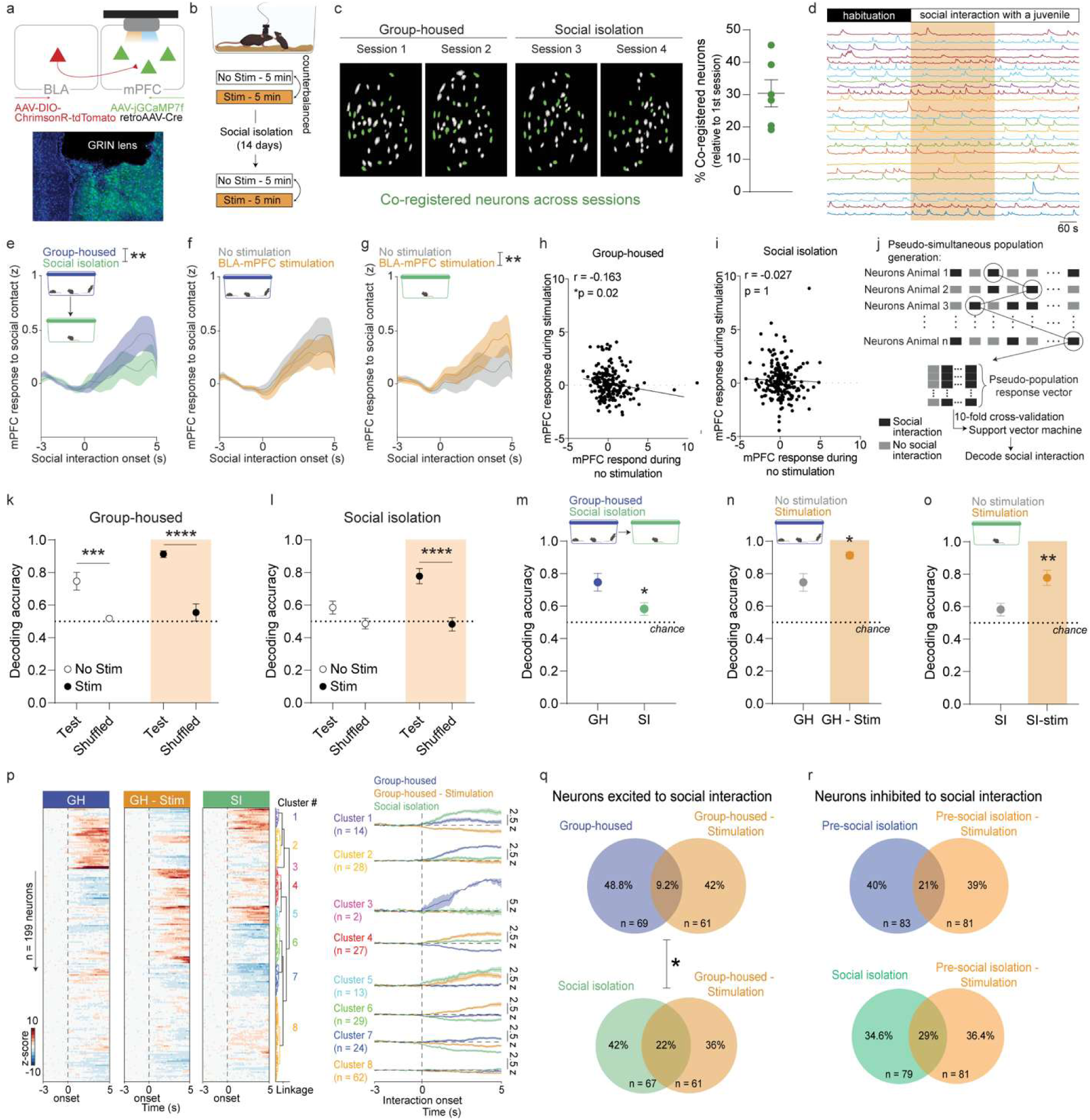
BLA-mPFC stimulation recruits a social isolation-like mPFC ensemble in response to social contact in males. **a.** Viral strategy and GRIN lens implant for mPFC imaging and BLA terminal stimulation, replicated in all mice. **b.** Recordings during the resident-juvenile intruder task before and after isolation; BLA-mPFC stimulation counterbalanced across days. **c. ∼**30% of mPFC neurons co-registered across all four sessions**. d.** Representative calcium traces during a stimulation session. **e.** Isolation decreases mPFC responses to interaction (*n* = 199 neurons/*N* = 6 male mice, two-way RM-ANOVA, interaction: F (49, 19404) = 1.538, \*\**p* = 0.009). **f.** BLA-mPFC stimulation did not affect mPFC responses before isolation (*n* = 199 co-registered neurons from *N* = 6 male mice). **g.** BLA-mPFC stimulation increases mPFC responses during isolation (*n* = 199 neurons/*N* = 6 male mice, two-way ANOVA, interaction: F (49, 7350) = 1.631, \*\**p* = 0.003). **h,i.** Stimulation and no-stimulation mPFC responses are negatively correlated in group-housed mice (h; *n* = 199 neurons/*N* = 6 male mice, Pearson correlation, *r*^2^ = 0.02, Bonferroni adjusted \**p* = 0.04) but uncorrelated during isolation (i; *r*^2^ = 0.0, *p* = 1). **j.** Pseudo-simultaneous population decoding schematic. **k.** mPFC decodes interaction above chance under both conditions (two-way ANOVA, main effect of group: F(1,36) = 54.29, \*\*\*\**p* < 0.0001; stimulation: F(1,36) = 6.47, \**p* = 0.01; Sidak’s post hoc, \*\*\**p* < 0.001). **l.** Isolation abolishes mPFC decoding, rescued by BLA-mPFC stimulation (two-way ANOVA, interaction: F (1, 36) = 5.95, \**p* < 0.05, main effect of group: F (1, 36) = 23.78, \*\*\*\**p* < 0.0001, stimulation: F (1, 36) = 5.47, \**p* = 0.01, Sidak’s post hoc, \*\*\*\**p* < 0.0001). **m.** Isolation decreases mPFC decoding accuracy (unpaired t-test, \**p* = 0.02). **n,o.** BLA-mPFC stimulation increases decoding accuracy in group-housed (n, unpaired t-test, \**p* = 0.01) and isolation (o, unpaired t-test, \**p* = 0.005) conditions. **p.** Functional mPFC clusters across group-housed and isolation sessions (*n* = 199 neurons/*N* = 6 male mice). **q.** BLA-mPFC stimulation recruits a mPFC ensemble resembling isolation-induced patterns, with greater overlap of excited neurons vs. group-housed (*n* = 199 neurons/*N* = 6 male mice*, Chi*^2^ = 6.94, \**p* = 0.03). **r.** Similar proportion of inhibited neurons recruited by stimulation in both conditions. Error bars indicate ±SEM. Schematics created in BioRender https://BioRender.com/qsp4lsv.

**Extended Data Figure 10.**
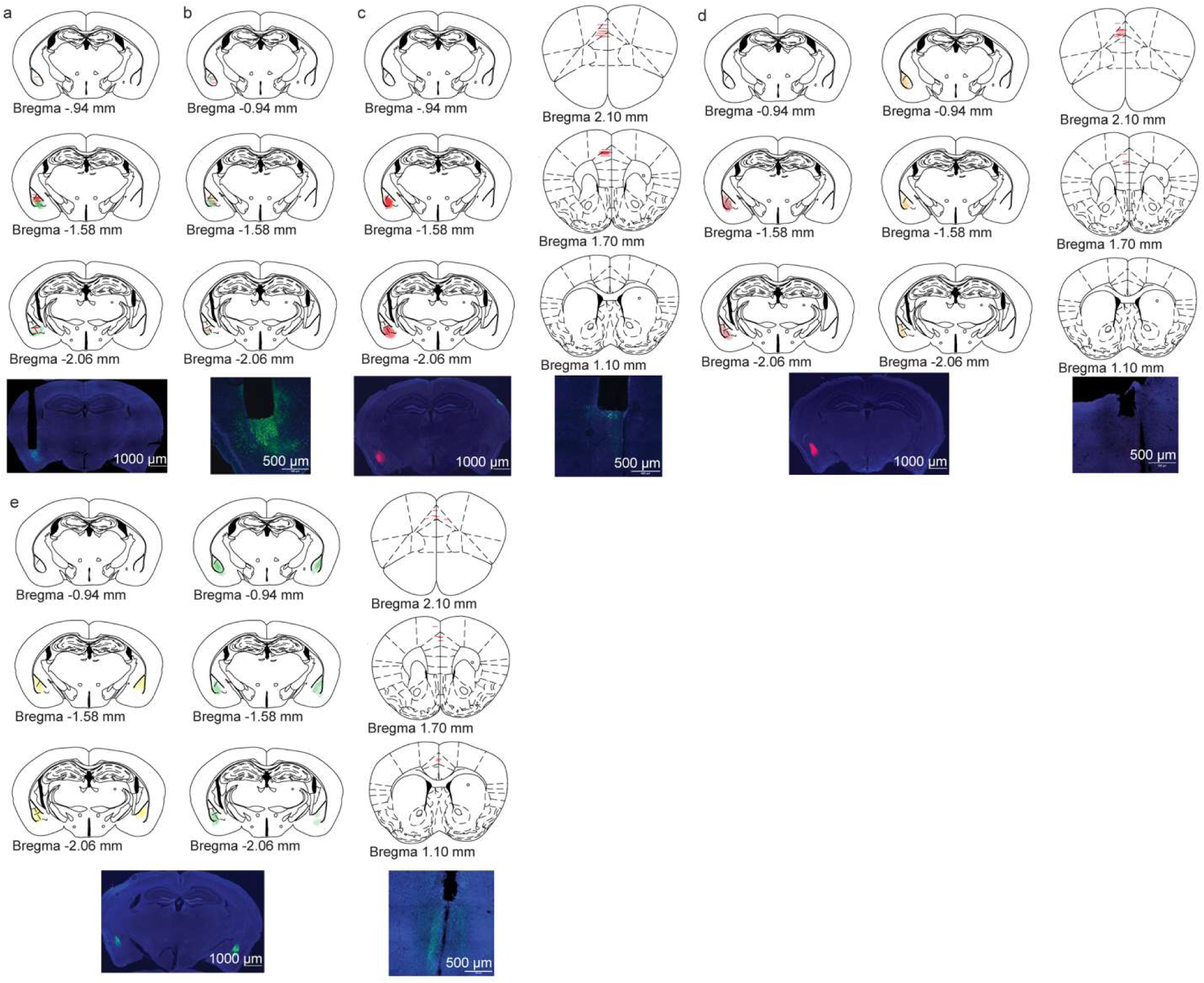
Histological verification of injection sites and implants. Injection sites for GCaMP7f and GRIN lens implant locations for non-specific BLA and BLA-mPFC calcium recordings, respectively. **c.** ChrimsonR expression in the BLA and GRIN lens implant locations in mPFC for simultaneous excitation of BLA-mPFC terminals and mPFC calcium recordings. **d.** ChrimsonR or tdTomato expression in BLA and fiber optic implant locations in mPFC for excitation of BLA-mPFC terminals during social interaction. **e.** NpHR or eYFP expression in the BLA and fiber optic implant locations in the mPFC for inhibition of BLA-mPFC terminals during social interaction and cued-2BC.

**Table S1.** Electrophysiological properties of BLA-mPFC and non-specific BLA neurons by housing condition. Unpaired t-test between pre-social isolation and social isolation conditions for each population, \**p* < 0.05.

| Sex | Condition | Population | Resting membrane potential (mV) | Capacitance (pF) | Input resistance (mΩ) | Threshold (pA) |
| --- | --- | --- | --- | --- | --- | --- |
| Male | GH | Non-specific BLA | -67.8 ± 0.63 | 164.2 ± 5.5 | 107.3 ± 4.0 | 175.5 ± 7.8 |
|  |  | BLA-mPFC | -68.4 ± 1.0 | 157.8 ± 8.0 | 98.6 ± 5.8 | 181.9 ± 12.7 |
| Male | SI | Non-specific BLA | -69.4 ± 0.5 | 170.9 ± 5.4 | 123.1 ± 5.8* | 146.4 ± 7.2** |
|  |  | BLA-mPFC | -69.1 ± 1.0 | 166.2 ± 8.0 | 118.4 ± 7.4* | 146.0 ± 14.3 |
| Female | GH | BLA-mPFC | -69.2 ± 1.0 | 126.3 ± 7.6 | 120.8 ± 7.5 | 97.9 ± 10.9 |
|  | SI | BLA-mPFC | -69.9 ± 0.6 | 156.4 ± 6.3** | 95.3 ± 5.6* | 136.0 ± 11.2* |

**Table S2.** Electrophysiological properties of BLA-mPFC and non-specific BLA neurons by social rank under group-housed conditions.

| Sex | Condition | Population | Rank | Resting membrane potential (mV) | Capacitance (pF) | Input resistance (MΩ) | Threshold (pA) |
| --- | --- | --- | --- | --- | --- | --- | --- |
| Female | GH | BLA-mPFC | Rank 1 | -69.4 ± 1.0 | 127.2 ± 12.6 | 110.2 ± 10.0 | 112.5 ± 16.0 |
| Female | GH | BLA-mPFC | Rank 2 | -66.1 ± 1.9 | 131.3 ± 11.6 | 116.8 ± 15.1 | 83.33 ± 24.9 |
| Female | GH | BLA-mPFC | Rank 3 | -73.6 ± 1.8 | 118.8 ± 18.1 | 142.8 ± 12.5 | 92.0 ± 14.9 |
| Male | GH | BLA-mPFC | Rank 1 | -67.3 ± 1.2 | 167.1 ± 10.5 | 95.1 ± 14.8 | 173.8 ± 22.4 |
| Male | GH | BLA-mPFC | Rank 2 | -69.0 ± 1.5 | 165.5 ± 15.9 | 99.5 ± 10.1 | 168.9 ± 20.5 |
| Male | GH | BLA-mPFC | Rank 3 | -69.3 ± 0.9 | 160.0 ± 9.5 | 91.55 ± 7.4 | 175.7 ± 12.2 |
| Male | GH | Non-specific BLA | Rank 1 | -70.9 ± 1.7 | 153.2 ± 8.5 | 113.0 ± 11.8 | 174.0 ± 24.6 |
| Male | GH | Non-specific BLA | Rank 2 | -63.2 ± 2.9 | 150.2 ± 15.1 | 124.2 ± 10.3 | 120.0 ± 17.3 |
| Male | GH | Non-specific BLA | Rank 3 | -66.7 ± 1.4 | 153.8 ± 9.6 | 105.7 ± 8.8 | 175.6 ± 16.2 |
| Male | SI | Non-specific BLA | Rank 1 | -69.4 ± 0.9 | 179.6 ± 11.7 | 113.5 ± 7.1 | 148.8 ± 13.1 |
| Male | SI | Non-specific BLA | Rank 2 | -70.2 ± 1.3 | 194.5 ± 19.6 | 115.8 ± 10.0 | 165.7 ± 17.6 |
| Male | SI | Non-specific BLA | Rank 3 | -69.3 ± 1.5 | 177.4 ± 15.5 | 99.7 ± 7.9 | 148.0 ± 20.5 |

